# Identification of distinct cDC2 subpopulations that direct microbiota-specific T cell differentiation

**DOI:** 10.1101/2025.11.04.686414

**Authors:** Shaina L. Carroll, Andrew Ly, Ashley K. Liu, Maria C. C. Canesso, Gabriel D. Victora, Daniel Mucida, Gregory M. Barton

**Affiliations:** Division of Immunology and Molecular Medicine, Department of Molecular and Cell Biology, University of California, Berkeley, CA, USA; Department of Plant and Microbial Biology, University of California, Berkeley, CA, USA; Laboratory of Mucosal Immunology, The Rockefeller University, New York, NY, USA; Laboratory of Lymphocyte Dynamics, The Rockefeller University, New York, NY, USA; Howard Hughes Medical Institute, The Rockefeller University, New York, NY, USA; Howard Hughes Medical Institute, University of California, Berkeley, CA, USA

**Author notes:** These authors contributed equally to this work (Shaina L. Carroll, Andrew Ly). Albert Einstein College of Medicine, Bronx, New York, USA.

## Abstract

How the complex network of intestinal antigen presenting cells (APCs) instructs CD4^+^ T cell responses against the microbiota remains unclear. Here, we use Labeling Immune Partnerships by SorTagging Intercellular Contacts (LIPSTIC) to characterize the APCs that prime CD4^+^ T cells recognizing the commensal bacterium *Akkermansia muciniphila*. *A. muciniphila*-specific T cells engaged multiple transcriptionally distinct migratory cDC2 subpopulations, both at homeostasis, when *A. muciniphil*a promotes T_FH_ differentiation, and during inflammation, when it also drives T_H_1 and T_H_17 differentiation. The identity of these subpopulations was unchanged by inflammation; however, the distribution of presentation across the subpopulations shifted, with increased presentation by inflammatory cDC2s favoring T_H_1 and T_H_17 polarization. These results reveal how distinct T cell differentiation trajectories can be determined through varied interactions with multiple, functionally distinct subpopulations of APCs.

## Introduction

The intestinal microbiota plays a fundamental role in shaping the host immune system. Of the hundreds of bacterial species that comprise the microbiota, only a handful have been identified that elicit antigen-specific CD4^+^ T cell responses (*1–5*). These anti-commensal T cells can contribute to tissue homeostasis and influence disease pathology in the context of both autoimmunity and cancer (*6–9*). Notably, each species known to elicit a CD4^+^ T cell response directs a unique T cell differentiation trajectory. For example, segmented filamentous bacteria (SFB) promote the differentiation of T_H_17 cells (*1, 10*), whereas *Helicobacter hepaticus* promotes the differentiation of regulatory T cells (T_regs_) (*5, 11*). What determines the nature of these commensal-specific responses remains poorly understood.

We have previously shown that *Akkermansia muciniphila,* a commensal bacterial species known to influence host physiology (*12–16*), induces antigen-specific CD4^+^ T cells (*4*). In gnotobiotic mice, this response is restricted to T follicular helper (T_FH_) cells in Peyer’s patches (PP), but in specific pathogen-free (SPF) mice, it expands to include T_H_1 and T_H_17 cells that traffic to the intestinal lamina propria (LP). The mechanisms underlying the variable T cell response elicited against *A. muciniphila* have not been examined.

A critical step that influences the differentiation of CD4^+^ T cells is their initial priming by antigen presenting cells (APCs) (*17*). During this process, APCs convey information to naive T cells about the nature of the presented antigen via surface-expressed ligands and secreted cytokines, guiding their differentiation into distinct effector subsets (*18*). The signals communicated by APCs depend on multiple factors, including the identity of the APC, environmental cues received from microbes, and the tissue microenvironment where priming occurs.

In the gut, APCs are diverse and include multiple conventional dendritic cell (cDC) subsets, monocytes, macrophages, and non-classical APCs, such as the recently described Rorγt^+^ APCs (*19–21*). Among these, cDCs have traditionally been considered the primary APC responsible for priming naive T cells. Broadly, cDCs can be divided into two major subsets, cDC1s and cDC2s (*22, 23*). In general, cDC1s specialize in priming T_H_1 responses as well as CD8^+^ T cells, whereas cDC2s have been shown to prime T_H_2, T_H_17, and T_FH_ responses (*24*). How cDC2s can prime such diverse effector fates is largely an open question. cDC2s are a heterogeneous population, and several recent studies, largely focused on splenic cDCs, have proposed a subdivision of cDC2s into cDC2As and cDC2Bs subsets, each with distinct phenotypes and functions. cDC2As display an anti-inflammatory profile, whereas cDC2Bs display a more pro-inflammatory program and are superior at promoting T_H_17 differentiation *in vitro* (*25, 26*). Therefore, one possible model is that distinct T cell effector subsets are instructed by distinct cDC2 subsets or states. However, this has been difficult to test experimentally due to the lack of tools for precisely depleting individual cDC2 subpopulations. The recently developed Labeling Immune Partnerships by SorTagging Intercellular Contacts (LIPSTIC) system overcomes this barrier by using a kiss-and-run labeling reaction to directly identify the APCs that engage with antigen-specific T cells during priming (*27, 28*).

In the intestine, the phenotype of APCs and CD4^+^ T cell responses are further influenced by the site of T cell priming. The gut-draining lymph nodes (gLNs) are anatomically and immunologically distinct; each gLN drains a defined segment of the gut (e.g., duodenum, jejunum, ileum, cecum and colon) and is characterized by a distinct immune tone (*29–31*). The immunologically distinct environment of each gLN can provide bystander signals that tune the functions of APCs and, consequently, the priming of CD4^+^ T cells. For example, the proximal gLNs that drain the duodenum and jejunum favor T_REG_ responses, while the distal gLNs that drain the ileum and large intestines favor inflammatory responses. However, this phenomenon has been described for only a few antigens, and it remains unclear how generalizable it is, particularly for commensal-derived antigens.

To better understand the mechanisms governing the differentiation of microbiota-reactive T cells, we combined the LIPSTIC system with single-cell RNA sequencing to characterize the APCs involved in priming *A. muciniphila*-specific T cells in diverse contexts where the responses differed. We demonstrate that *A. muciniphila*-specific T cells are primed by multiple transcriptionally distinct subpopulations of migratory cDC2s, both at homeostasis and during inflammation. Surprisingly, we find that the identities of these cDC2s do not change between the two contexts. Instead, inflammation alters the relative abundance of the subpopulations presenting *A. muciniphila*-presenting antigen, with a shift toward inflammatory cDC2s that drive pro-inflammatory fates in *A. muciniphila*-specific T cells.

## Results

### A. muciniphila-specific T cells are activated in both the gut-draining lymph nodes and the Peyer’s patches

We have previously shown that *A. muciniphila*-specific T cells adopt a T_FH_ phenotype in the PPs of mice colonized with a minimal eight-member microbiota known as Altered Schaedler’s flora (ASF) along with an *A. muciniphila* strain previously isolated from our mouse colony (hereafter referred to as ASF+Akk mice) (*4*). However, it was unclear whether these cells were primed locally in the PPs or were recruited following priming at another site. To determine the relevant location(s) of *A. muciniphila*-specific T cell priming, we tracked the activation of naive *A. muciniphila*-specific Am124 CD4^+^ T cells (with a Thy1.1 congenic marker) over the course of a week (Fig. 1A). We directed our analysis to the gLNs and PPs, the two main inductive sites for adaptive immune responses in the gut (*32*). Am124 T cells accumulated in both tissue sites starting five days post-transfer (Fig. 1, B and C). However, Am124 T cells upregulated the T-cell activation marker CD69 as early as one day post-transfer with a peak at two days post-transfer in both tissues, indicating that T cells are being activated within 48 hours (Fig. 1, D and E). Am124 T cells also upregulated Bcl6, the T_FH_-lineage-defining transcription factor, as early as one day post-transfer in the gLNs (Fig. 1F). In the PPs, Bcl6 expression was delayed and was only consistently detected in Am124 T cells five days post-transfer (Fig. 1G). Altogether, these data indicate that the priming and initial polarization of *A. muciniphila*-specific T_FH_ cells occurs in both the gLNs and the PPs at steady state.

**Figure 1.**
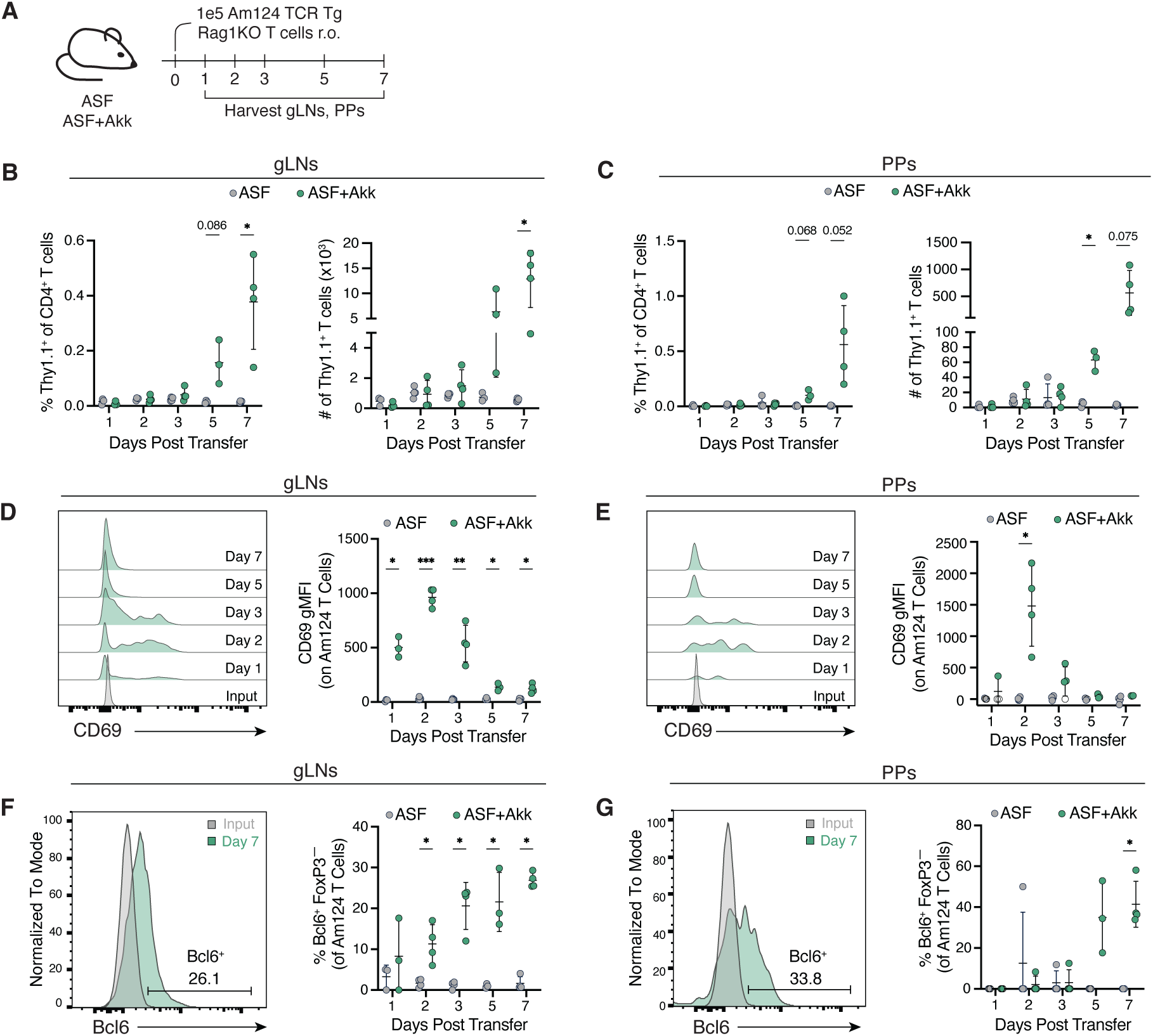
*A. mucinip*hila-specific T cells are activated in both the gLNs and the Peyer’s patches. **(A)** Schematic representation of experimental timeline. 1×10^5^ Am124 T cells were adoptively transferred into ASF and ASF+Akk mice. gLNs and PPs were harvested 1, 2, 3, 5, and 7 days post-transfer. (**B** and **C**) Frequency (left) and numbers (right) of *A. muciniphila*-specific T cells at days 1, 2, 3, 5 and 7 post transfer in the gLNs (**B**) and PPs (**C**). (**D** and **E**) Representative histograms (left) and quantification (right) of CD69 expression on Am124 T cells at days 0 (input), 1, 2, 3, 5 and 7 post transfer in the gLNs (**D**) and PPs (**E**). (**F** and **G**) Representative histograms (left) and quantification (right) of Bcl6 expression on *A. muciniphila*-specific T cells at days 0 (input), 1, 2, 3, 5 and 7 post transfer in the gLNs (**F**) and PPs (**G**). For all graphs, each symbol represents one mouse and error bars represent mean and standard deviation (Am124 T cells were not found in the PPs of some mice; these mice are represented by an open circle in (E)); n = 3 to 4 mice per group, data is representative of two independent experiments. P-values were calculated via unpaired T-test with Welch correction. Statistical significance denoted as *P < 0.05, **P < 0.01, ***P < 0.001, ****P < 0.0001.

### A. muciniphila- and SFB-specific T cells are activated in the same gut-draining lymph nodes but differentiate into distinct T helper types

After identifying the location where *A. muciniphila*-specific T cells are primed, we next aimed to understand the cues driving the differentiation of these cells towards particular effector phenotypes. One possible explanation for the biased T_FH_ response to *A. muciniphila* in ASF mice is that the ASF microbiota lacks a strong T_H_17 inducer (*33*), in contrast to our conventionally-housed SPF mice, which are typically colonized with SFB and in which we previously reported mixed *A. muciniphila*-specific T cell phenotypes(*4*). Therefore, to test whether strong T_H_17 polarizing signals can influence *A. muciniphila*-specific T cell differentiation, we colonized ASF+Akk mice with SFB and examined the differentiation of both Am124 T cells (Thy1.1^+^) and SFB-specific 7B8 T cells (CD45.1^+^) in these ASF+Akk+SFB mice (Fig. 2A).

**Figure 2.**
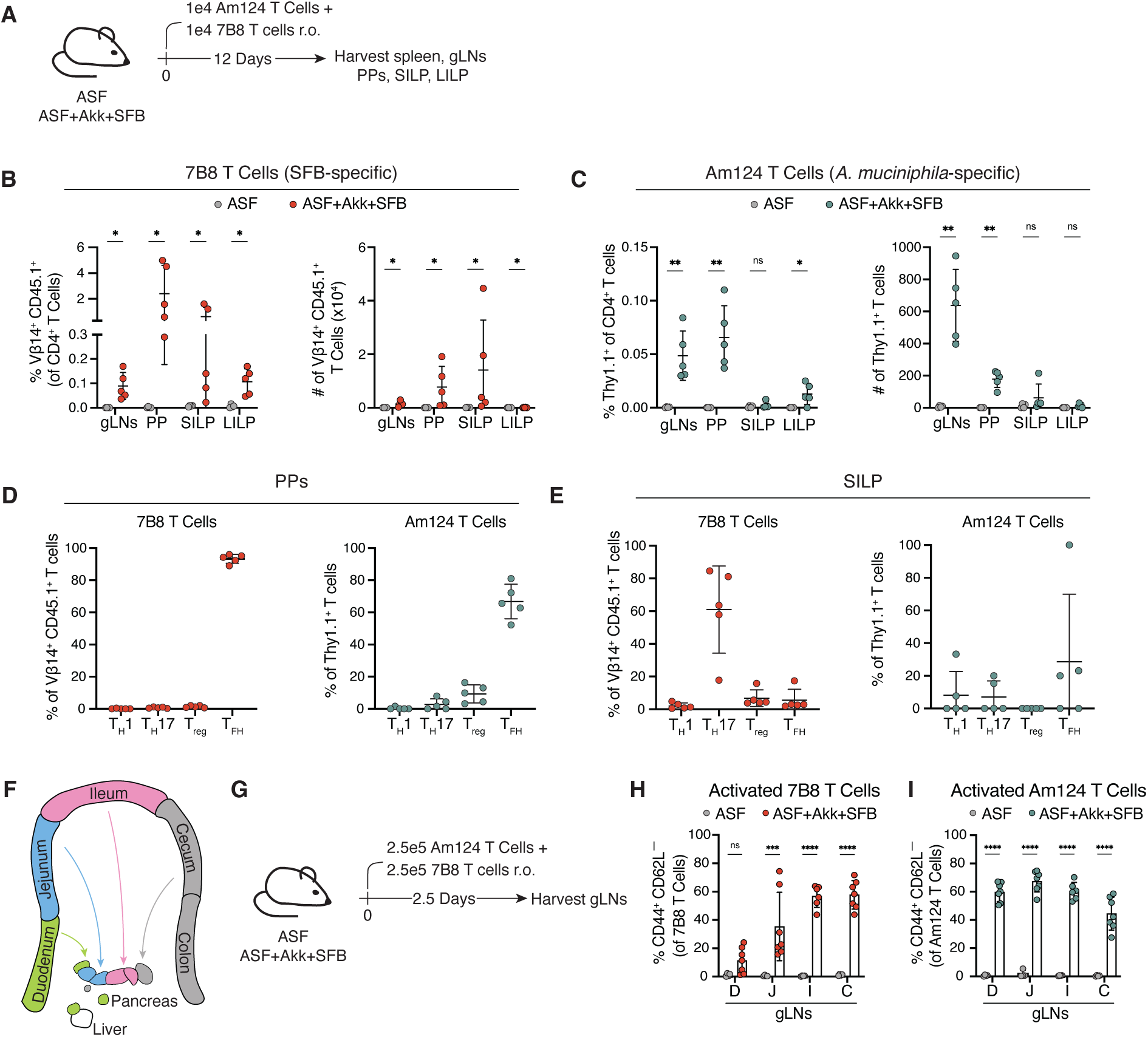
*A. muciniphila*- and SFB-specific T cells are activated in the same gLNs but differentiate into distinct T helper types. (**A**) Schematic representation of experimental timeline. 1×10^4^ Am124 and 1×10^4^ 7B8 T cells were adoptively transferred simultaneously into ASF and ASF+Akk+SFB mice. gLNs, PPs, SILP, and LILP were harvested 12 days later. (**B**) Frequency (left) and numbers (right) of transferred 7B8 T cells in tissues of ASF and ASF+Akk+SFB mice 12 days post transfer. (**C**) Frequency (left) and numbers (right) of transferred Am124 T cells in tissues of ASF and ASF+Akk+SFB mice 12 days post transfer. (**D**) Expression of T_H_1 (Tbet^+^ FoxP3^−^), T_H_17 (Rorγt^+^ FoxP3^−^), T_REG_ (FoxP3^+^), and T_FH_ (FoxP3^−^ Bcl6^+^ PD-1^+^) markers in transferred 7B8 (left) and Am124 (right) T cells in the PPs of ASF+Akk+SFB mice. (**E**) Expression of T_H_1 (Tbet^+^ FoxP3^−^), T_H_17 (Rorγt^+^ FoxP3^−^), T_REG_ (FoxP3^+^), and T_FH_ (FoxP3^−^ Bcl6^+^ PD-1^+^) markers in transferred 7B8 (left) and Am124 (right) T cells in the SILP of ASF+Akk+SFB mice. For (**B** to **E**), n = 4 to 5 mice per group, data is representative of three independent experiments. (**F**) Diagram of segmented gLNs harvested for analysis. (**G**) Schematic representation of experimental timeline for (**H** and **I**). 2.5×10^5^ Am124 and 2.5×10^5^ 7B8 T cells were adoptively transferred simultaneously into ASF and ASF+Akk+SFB mice and gLNs were harvested and segmented based on drainage 2.5 days later. (**H**) Frequency of activated (CD44^+^ CD62L^−^) 7B8 T cells in gLNs. (**I**) Frequency of activated (CD44^+^ CD62L^−^) Am124 T cells in gLNs. For (**H** and **I**), n = 4 to 6 mice per group; data is representative of two independent experiments. D, duodenum; J, jejunum; I, ileum; C, cecum and ascending colon. For all graphs, each symbol represents one mouse and error bars represent mean and standard deviation. P-values were calculated by Mann-Whitney test for (B); by unpaired T-test with Welch correction for (C); and by two-way ANOVA for (H and I). Statistical significance denoted as not significant (ns), *P < 0.05, **P < 0.01, ***P < 0.001, ****P < 0.0001.

In line with previously published data, 7B8 T cells in ASF+Akk+SFB mice localized to the small and large intestine lamina propria (SILP and LILP), in addition to the PPs and gLNs (*1, 34*) (Fig. 2B). In contrast, Am124 T cells trafficked mainly to the gLNs and PPs, with few cells detected in the SILP and LILP (Fig. 2C). In the PPs, both Am124 and 7B8 T cells adopted T_FH_ phenotypes (Bcl6^+^ PD-1^+^ FoxP3^−^) (Fig. 2D, and fig. S1A). In the SILP, 7B8 T cells primarily differentiated into T_H_17 cells (Rorγt^+^ FoxP3^−^), whereas the very few Am124 T cells found in the SILP did not exhibit strong polarization towards one specific CD4^+^ T cell fate (Fig. 2E, and fig. S1B). *A. muciniphila*- and SFB-specific CD4^+^ T cells thus adopt distinct T_H_ identities and tissue localization patterns in ASF+Akk+SFB mice. Importantly, the phenotype and trafficking patterns of Am124 T cells in ASF+Akk+SFB mice are consistent with what we have previously described in ASF+Akk mice (*4*), indicating that the differentiation of these cells is not influenced by the presence of a strong T_H_17-polarizing microbe, such as SFB.

One possibility for the distinct differentiation of 7B8 and Am124 T cells in ASF+Akk+SFB mice is that these cells are primed in different gLNs, each possessing unique immunological tones that promote distinct CD4^+^ T cell differentiation trajectories (*29–31*). To address this hypothesis, we co-transferred naive 7B8 and Am124 CD4+ T cells into ASF+Akk+SFB mice and assessed their priming in segmented gLNs 2.5 days post-transfer, a timepoint at which we observe peak activation of Am124 T cells in the gLNs (Fig. 1D and Fig. 2, F and G). Consistent with prior findings, we observed that 7B8 T cells upregulated the T cell activation marker CD44, mainly in the ileal and cecal-colonic gLNs (*30*) (Fig. 2H, and fig. S1C). In contrast, Am124 T cells upregulated CD44 in all the segmented gLNs analyzed (Fig. 2I, and fig. S1D). Thus, *A. muciniphila*- and SFB-specific CD4^+^ T cells are primed within the same gLNs, despite differentiating into distinct T_H_ fates, indicating that the unique immune tone of individual gLNs does not fully explain the divergence of T_H_ differentiation programs among these commensal-specific CD4^+^ T cells.

### A. muciniphila is presented by a transcriptionally distinct population of migratory cDC2s at steady state

Our previous results indicate that neither the absence of a strong T_H_17-polarizing microbe in the ASF microbiota nor the site of T cell priming in the gLNs is sufficient to explain the unique T_FH_-dominated response elicited against *A. muciniphila* in ASF+Akk animals. Thus, we next sought to identify and characterize the APCs that direct *A. muciniphila*-specific CD4^+^ T cells towards their distinct differentiation trajectory at steady state. To identify cells presenting *A. muciniphila* antigens, we used the LIPSTIC system (*27*). This system relies on the transpeptidase sortase A (SrtA), which catalyzes the transfer of LPXTG peptides to nearby strings of N-terminal oligoglycines. In the LIPSTIC system, a CD40L-SrtA fusion protein expressed on recently activated T cells (*35*) transfers LPETG-biotin to a string of five glycines expressed at the extracellular N-terminus of CD40 on interacting APCs (Fig. 3A). The labeled (biotin^+^) cognate APCs can then be detected using an anti-biotin antibody. Although LIPSTIC has previously been used to investigate APC–T cell interactions in the intestine in the context of oral tolerance using the model antigen ovalbumin (*36*), it has not yet been used to characterize T cell responses against commensal microbes.

**Figure 3:**
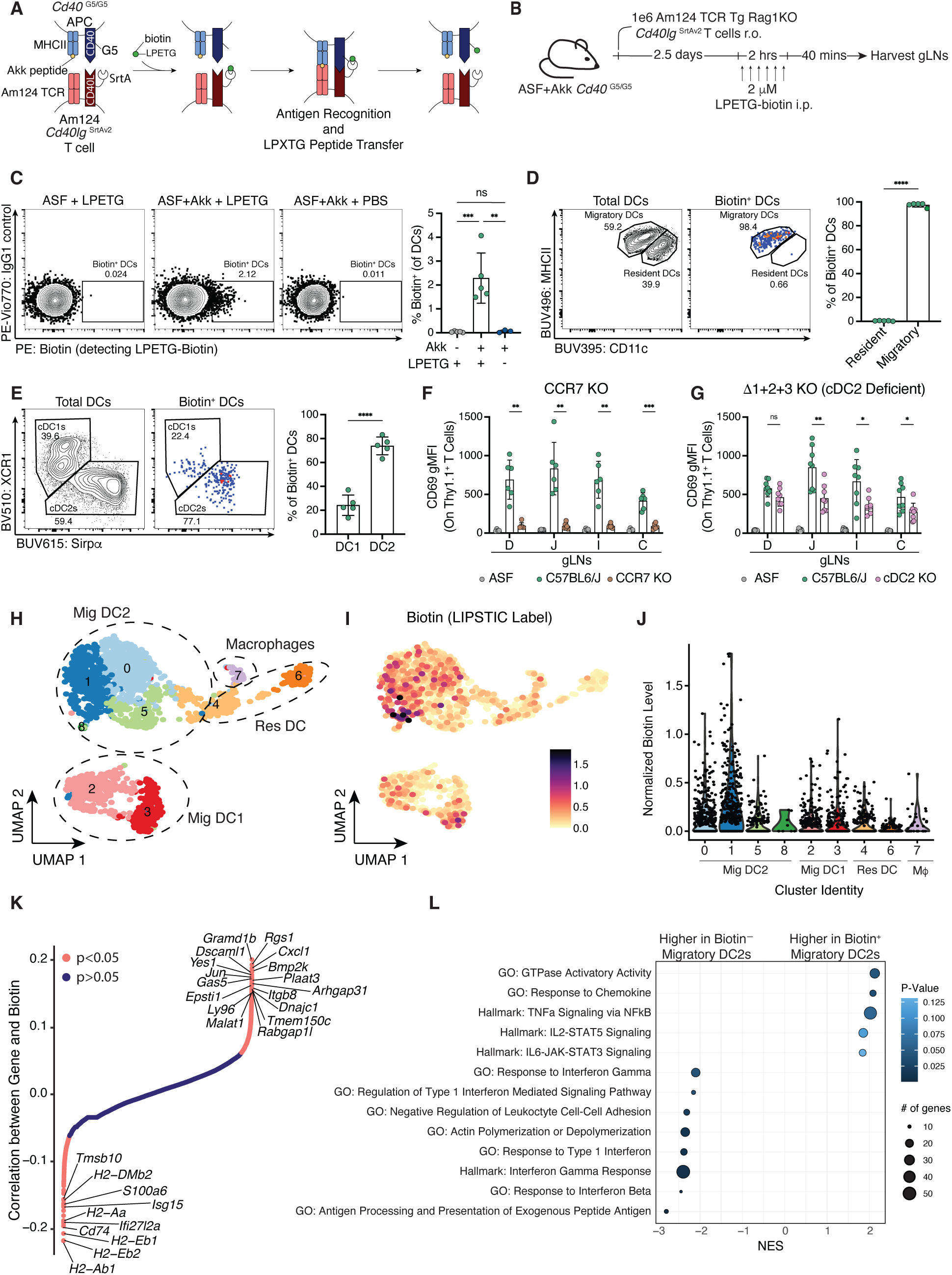
***A. muciniphila* is presented by a transcriptionally distinct population of migratory cDC2s at steady state.** (**A**) Schematic representation of LIPSTIC labeling with *Cd40lg*^SrtAv2^ Am124 TCR transgenic T cells. (**B**) Experimental timeline for LIPSTIC labeling experiments. 1×10^6^ *Cd40lg*^SrtAv2^ Am124 T cells were adoptively transferred into ASF or ASF+Akk mice and LIPSTIC labeling was performed 2.5 days later to identify *A. muciniphila*-presenting cells. (**C**) Representative flow plots (left) and quantification (right) showing percentage of labeled cDCs in gLNs. (**D**) Representative flow plots showing gating for resident and migratory cDCs on total cDCs (left) and biotin^+^ cDCs (center) and quantification of data (right). (**E**) Representative flow plots showing gating for cDC1s (Xcr1^+^) and cDC2s (Sirpα^+^) on total cDCs (left) and biotin^+^ cDCs (center) and quantification of data (right). For (**C** to **E**), n = 3 to 5 mice per group; data is representative of three independent experiments. For (**F** and **G**), 1.5×10^5^Am124 T cells were adoptively transferred into ASF control and ASF+Akk BMC mice of the indicated genotype and gLNs were harvested 1.5 days later. (**F**) Expression of CD69 on Am124 T cells in gLNs of ASF control mice and ASF+Akk BMCs reconstituted with C57BL/6J or *Ccr7^−^*^/–^ bone marrow (n = 4 to 6 mice per group, data is representative of two independent experiments). (**G**) Expression of CD69 on Am124 T cells in gLNs of ASF control mice and ASF+Akk BMCs reconstituted with C57BL/6J or Δ1+2+3 bone marrow (n = 3 to 4 mice per group, per experiment, data is pooled from two independent experiments). For (**H** to **L**), 1×10^6^ *Cd40lg*^SrtAv2^ Am124 T cells were adoptively transferred into ASF+Akk mice (n = 5). LIPSTIC labeling was performed 2.5 days later to sort and sequence *A. muciniphila*-presenting cells. (**H**) Uniform manifold approximation and projection (UMAP) plot of sorted and sequenced DCs from gLNs of ASF+Akk mice. Cells are grouped into clusters via unsupervised hierarchical clustering. Circles indicating cDC subset were drawn based on expression of key marker genes as shown in fig. S5. (**I**) Log normalized counts of LIPSTIC signal in the same cells from (H) detected via anti-PE hashtag antibody. (**J**) Violin plot showing log normalized counts of LIPSTIC signal in each cluster. Each dot represents one cell. (**K**) Spearman correlation between scaled gene expression and normalized LIPSTIC signal, calculated for all genes in migratory cDC2s. Each gene is plotted in order of increasing correlation with LIPSTIC signal. (**I**) Pathway analysis for genes that were significantly positively or negatively correlated with the LIPSTIC signal in (K). Size of the dot indicates how many genes from each gene signature were present among the significantly correlated genes. Color of the dot indicates p-values. For (C to G), each symbol represents one mouse and error bars represent mean and standard deviation. P-values were calculated by one-way ANOVA for (C); unpaired T-test for (D), and (E); unpaired T-test with Welch’s correction for (F), and (G); and by permutation test with correction for multiple hypothesis test using the BH method for (K). Statistical significance denoted as not significant (ns), *P < 0.05, **P < 0.01, ***P < 0.001, ****P < 0.0001.

To apply the LIPSTIC system in the context of the ASF microbiota, we adoptively transferred Am124 *Cd40lg*^SrtAv2^ T cells into adult ASF or ASF+Akk *Cd40*^G5/G5^ recipients and performed LIPSTIC labeling via intraperitoneal injection of the LPETG-biotin substrate 2.5 days later (Fig. 3B). Due to the limited number of Am124 T cells found in the PPs at this timepoint, we examined labeling of APCs in the gLNs only (Fig. 1C). Using this approach, we observed that approximately 2% of cDCs were labeled with biotin in mice colonized with *A. muciniphila* that received the LPETG substrate, while no labeling was observed in control mice either lacking *A. muciniphila* or not receiving the LPETG substrate (Fig. 3C, and fig. S2A). Previous work has shown that LIPSTIC labeling at late timepoints can result from non-cognate interactions between CD4^+^ T cells and bystander cDCs (*27*). To confirm that the labeling we observed at this 2.5 day timepoint was due to cognate interactions, we generated mixed bone marrow chimeras (BMCs) in which 25% of cDCs lack MHCII expression (*H2-Ab1^-/-^*). All cDCs in these mice still expressed the *Cd40*^G5/G5^ allele, allowing them to be labeled. To maintain gnotobiotic conditions in these chimeras, we used busulfan rather than traditional irradiation for host conditioning fig. S2B) (*35*). As expected, labeling occurred only on MHCII^+^ cDCs in these mixed BMCs, confirming that the observed labeling required cognate MHCII-TCR interactions (fig. S2, C to E).

Biotin^+^ cDCs were uniformly migratory cDCs (MHCII^hi^ CD11c^int^); we observed no labeling of LN-resident cDCs (MHCII^int^CD11c^hi^) in the gLNs (Fig. 3D). This selective labeling of migratory cDCs aligns with prior LIPSTIC studies showing that T cells also exclusively interact with migratory cDCs in the settings of immunization, oral tolerance, and immune checkpoint blockade (*27, 36, 37*). The majority of biotin^+^ cDCs were CD103^+^ cDC2s (Sirpɑ^+^ CD11b^+^), although we did consistently observe a small population of biotin^+^ cDC1s (Xcr1^+^) (Fig. 3E and fig. S2, F and G). We also performed LIPSTIC labeling in mice colonized with more complex microbiota, including the 12-member OligoMouse Microbiota (OMM^12^) (*38*) and an SPF microbiota. In OMM^12^ and SPF *Cd40^G5/G5^*mice, migratory cDC2s remained the predominant interacting APC (fig. S3, A to C, and fig. S3, F to H). Although we have previously reported that Am124 T cells can differentiate into multiple effector phenotypes in conventionally-housed SPF mice (*4*), we did not observe the same diverse response in SPF mice housed in a high-barrier facility. In these barrier SPF mice, as well as in OMM^12^ mice, Am124 T cells primarily differentiated into T_FH_ cells in Peyer’s patches without the appearance of additional T_H_ subsets (fig. S3, D and E and fig. S3, I and J).

To genetically confirm that migratory cDC2s are required for priming of *A. muciniphila-*specific T cells, we again turned to the busulfan BMC system. Transferred Am124 T cells failed to upregulate CD69 in *Zbtb46*-DTR chimeras depleted of cDCs (*39*), confirming the dominant role for cDCs in priming *A. muciniphila*-specific T cells (fig. S4, A and B). Similarly, in *Ccr7*^-/-^ BMCs, in which hematopoietic cells are unable to migrate from the LN (*39*), transferred Am124 T cells also failed to upregulate CD69 in the gLNs, indicating that *A. muciniphila* antigens must be actively carried to the gLNs by migratory cells for presentation to T cells (Fig. 3F, and fig. S4, C to E). In BMCs with reduced cDC2s (generated using *Zeb2* triple enhancer mutant mice, known as Δ1+2+3 mice, in which mature cDC2s fail to develop from myeloid progenitors (*40*)), there was also reduced upregulation of CD69 on transferred Am124 T cells in the gLNs (Fig. 3G and fig. S4H). Although we consistently observed a small population of biotin^+^ cDC1s, we found that cDC1s play a negligible role in priming Am124 T cells, as there was no defect in CD69 upregulation on transferred Am124 T cells in the gLNs of *Batf3*^-/-^ BMCs, lacking cDC1s (*41*) (fig. S4, F and G). Taken together with the LIPSTIC labeling experiments, these results show that *A. muciniphila* is primarily presented by migratory cDC2s in the gLNs at steady state when *A. mucinphila* T cells mainly adopt a T_FH_ phenotype. While previous studies have implicated cDC2s in priming T_FH_ responses to non-commensal antigens, our findings demonstrate for the first time that cDC2s are essential for the priming of T_FH_ responses against commensal antigens (*42–45*).

As noted earlier, cDC2s are a heterogeneous population of cells comprising several subpopulations. Therefore, we hypothesized that *A. muciniphila*-specific T cells might be primed by a distinct population of cDC2s with a T_FH_-inducing ability. To identify such a subpopulation, we performed single-cell RNA sequencing (scRNA-seq) on biotin^−^ and biotin^+^ cDCs sorted from the gLNs of LIPSTIC-labeled mice (fig. S5A). We included CITE-seq antibodies (*46*) against PE (to detect the anti-biotin PE), as well as key cDC subset markers (CD11b, XCR1) (fig. S5B).

Clustering analysis revealed eight transcriptionally distinct cDC populations and one macrophage cluster (cluster 7), with all clusters represented across all mice (Fig. 3H, and fig. S5, C and D). Clusters 2 and 3 were identified as migratory cDC1s based on expression of XCR1 protein and *Ccr7* transcript. Clusters 0, 1, 5, and 8 corresponded to migratory cDC2s, expressing CD11b protein and *Sirpɑ* and *Ccr7* transcripts. Cluster 6 represented resident cDC1s (high XCR1 protein, low *Ccr7* RNA), and cluster 4 included a mix of resident and migratory cDC2s, distinguishable from one another by *Ccr7* expression (fig. S5C). In line with the flow cytometry analysis, biotinylation was highest among the migratory cDC2 clusters in all mice (Fig. 3I and J, and fig. S5E). Interestingly, this signal was not uniform across all migratory cDC2 clusters: clusters 0 and 1 were the most highly biotinylated, whereas clusters 5, 8, and 4 had much lower levels of biotinylation.

To identify transcriptional programs that were differentially regulated between biotin^+^ and biotin^−^ migratory cDC2s, we performed a correlation analysis, taking advantage of the continuous nature of the biotin signal. We identified 545 genes that were positively correlated (enriched in biotin^+^ migratory cDC2s) and 496 genes that were negatively correlated with the biotin signal (enriched in biotin^−^ migratory cDC2s) (Fig. 3K, and Table S3). We then performed gene set enrichment analysis of these genes to identify pathways positively and negatively correlated with the biotin signal. Biotin^+^ migratory cDC2s scored highly for GTPase activation, response to chemokines, and NF-κB signaling (Fig. 3L). There was also a trend towards higher Stat5-IL2 signaling and Stat3-IL6 signaling in biotin^+^ cDCs, although these pathways did not reach statistical significance. Biotin^−^ cDCs scored highly for antigen processing and presentation, as well as both type 1 and type 2 interferon signaling. Thus, through a novel application of the LIPSTIC system to characterize commensal-specific T cell-APC interactions, we identified a distinct subpopulation of migratory cDC2s that present *A. muciniphila* at steady state. These cells display a transcriptional signature indicative of elevated activation, migratory potential, and reduced antigen processing/presentation, likely due to recent stimulation by *A. muciniphila* products during antigen acquisition.

### A. muciniphila-specific T cells adopt alternative fates during inflammation

Next, we sought to define how the distinct transcriptional signature of *A. muciniphila*-presenting cDCs related to the unique differentiation trajectory of *A. muciniphila*-specific CD4^+^ T cells. We reasoned that by comparing this signature to that of APCs directing different T cell fates, we would reveal critical determinants that drive each T_H_ type. We therefore worked to identify a context in which the *A. muciniphila*-specific T cell response is altered in ASF+Akk mice. Several genome-wide association studies have linked mutations in the IL-10 signaling pathway with inflammatory bowel diseases in humans (*47–51*), and previous studies in mice have shown that blocking IL-10 signaling can alter the differentiation of *H. hepaticus*-specific T cells from T_REG_ cells to T_H_17 cells (*5, 11*). To determine whether inflammation induced by IL-10 signaling blockade could skew the differentiation of *A. muciniphila*-specific T cells, we treated ASF+Akk mice with blocking antibodies against the IL-10 receptor. Three days after the initial antibody injection, we transferred Am124 T cells and examined their phenotype 12 days later (Fig. 4A). In ɑIL-10R-treated mice, a greater number of Am124 T cells accumulated in the SILP and LILP (Fig. 4, B and C), where they were more likely to differentiate into inflammatory T_H_1 and T_H_17 cells (Fig. 4, D-G, and fig. S6, A and B). Endogenous T cells in the LP of ɑIL-10R-treated mice also significantly upregulated Rorγt but not Tbet (fig. S6C). Treatment with ɑIL-10R did not alter the differentiation of Am124 T cells in the PPs, where they still adopted T_FH_ markers, nor did it induce large changes in the endogenous T cells or Am124 T cells in the gLNs (fig. S6D). This effect of ɑIL-10R was not due to changes in the distribution of *A. muciniphila*-specific T cell priming across the various gLNs or in the type of APC presenting *A. muciniphila,* as Am124 T cell activation across all segments of the gLNs was comparable between control and antibody-treated mice and LIPSTIC labeling revealed that the majority of biotin^+^ cells were migratory cDC2s in both control and antibody-treated mice (fig. S7, A to D). Overall, these results indicate that ɑIL-10R antibody treatment of ASF+Akk mice promotes the trafficking of *A. muciniphila*-specific T cells to the LPs and their differentiation into pro-inflammatory phenotypes in these tissues, without changing the site of priming or the type of APC presenting *A. muciniphila*.

**Figure 4:**
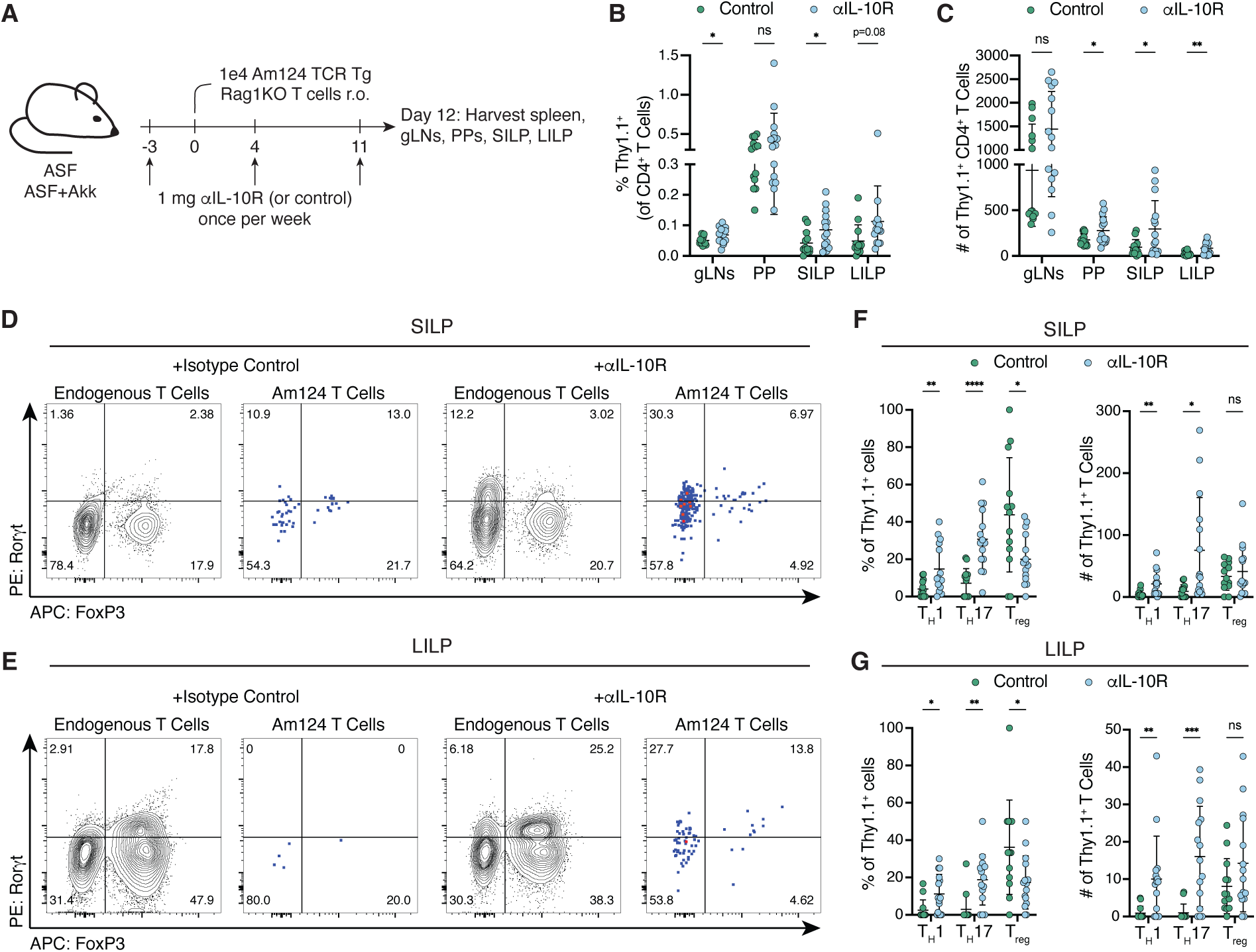
***A. muciniphila*-specific T cells adopt alternative fates during inflammation.** (**A**) Experimental timeline for (**B** to **G**). ASF+Akk mice were injected with 1 mg ɑIL-10R or PBS/isotype control antibodies once per week. 1×10^4^ Am124 T cells were adoptively transferred 3 days after the initial antibody injection and tissues were harvested 12 days after T cell transfer. (**B** and **C**) Frequency (**B**) and number (**C**) of transferred Am124 T cells in tissues of control and ɑIL-10R treated mice 12 days post transfer. (**D** and **E**) Representative flow plots showing expression of T_H_17 markers (Rorγt^+^ FoxP3^−^) in endogenous and transferred T cells in the SILP (**D**) and LILP (**E**) of control and ɑIL-10R treated mice. **(F** and **G)** Frequency (left) and number (right) of transferred Am124 T cells expressing T_H_1 (Tbet^+^ FoxP3^−^), T_H_17 (Rorγt^+^ FoxP3^−^), and T_REG_ (FoxP3^+^) markers in the SILP (**F**) and LILP (**G**) of control and ɑIL-10R-treated mice. For all graphs, each symbol represents one mouse and error bars represent mean and standard deviation; n = 4 to 6 mice per group, per experiment, data is pooled from three independent experiments. P-values were calculated via unpaired T-test. Statistical significance denoted as not significant (ns), *P < 0.05, **P < 0.01, ***P < 0.001, ****P < 0.0001.

### Inflammatory migratory cDC2s drive skewing of A. muciniphila-specific T cells towards alternative fates during inflammation

Based on these results, we considered that the shift in Am124 T cell differentiation was driven by changes in the phenotype of the presenting cDCs. To test this, we performed scRNA-seq on sorted cDCs from LIPSTIC-labeled mice treated with either isotype control or ɑIL-10R antibodies. In this experiment, we included a control sample from an ASF mouse lacking *A. muciniphila* enabling us to define the border between biotin^+^ and biotin^−^ cells (fig. S8B).

Clustering analysis revealed similar transcriptional clusters to what we had observed previously– several distinct clusters of resident cDCs, migratory cDC1s, and migratory cDC2s (Fig. 5A and fig. S8A). Surprisingly, biotin^−^ cDCs from ɑIL-10R-treated mice exhibited few transcriptional differences from controls (fig. S8D). All clusters contained cells from both treatment groups, indicating that blocking IL-10 signaling does not induce a distinct cDC subpopulation (fig. S8C). Among biotin^+^ DCs, most cells fell into three transcriptional clusters (clusters 1, 2, and 7), both at steady state and during inflammation, corresponding to migratory cDC1s (cluster 1) and migratory cDC2s (clusters 2 and 7) (Fig. 5, B and C, and fig. S8A). This is consistent with our flow cytometry data showing that *A. muciniphila*-presenting cells, both at steady state and during inflammation, are predominantly migratory cDC2s, with a smaller population of labeled migratory cDC1s. Since cDC1s are dispensable for the priming of Am124 T cells (fig. S4F) and the location of biotin^+^ cDC1s on the UMAP was the same in control and ɑIL-10R-treated mice (Fig. 5B), we concluded that transcriptional changes in *A. muciniphila*-presenting cDC1s were likely not responsible for the shift in the phenotype of Am124 T cells. Thus, we focused our subsequent analysis on *A. muciniphila*-presenting cDC2s.

**Figure 5:**
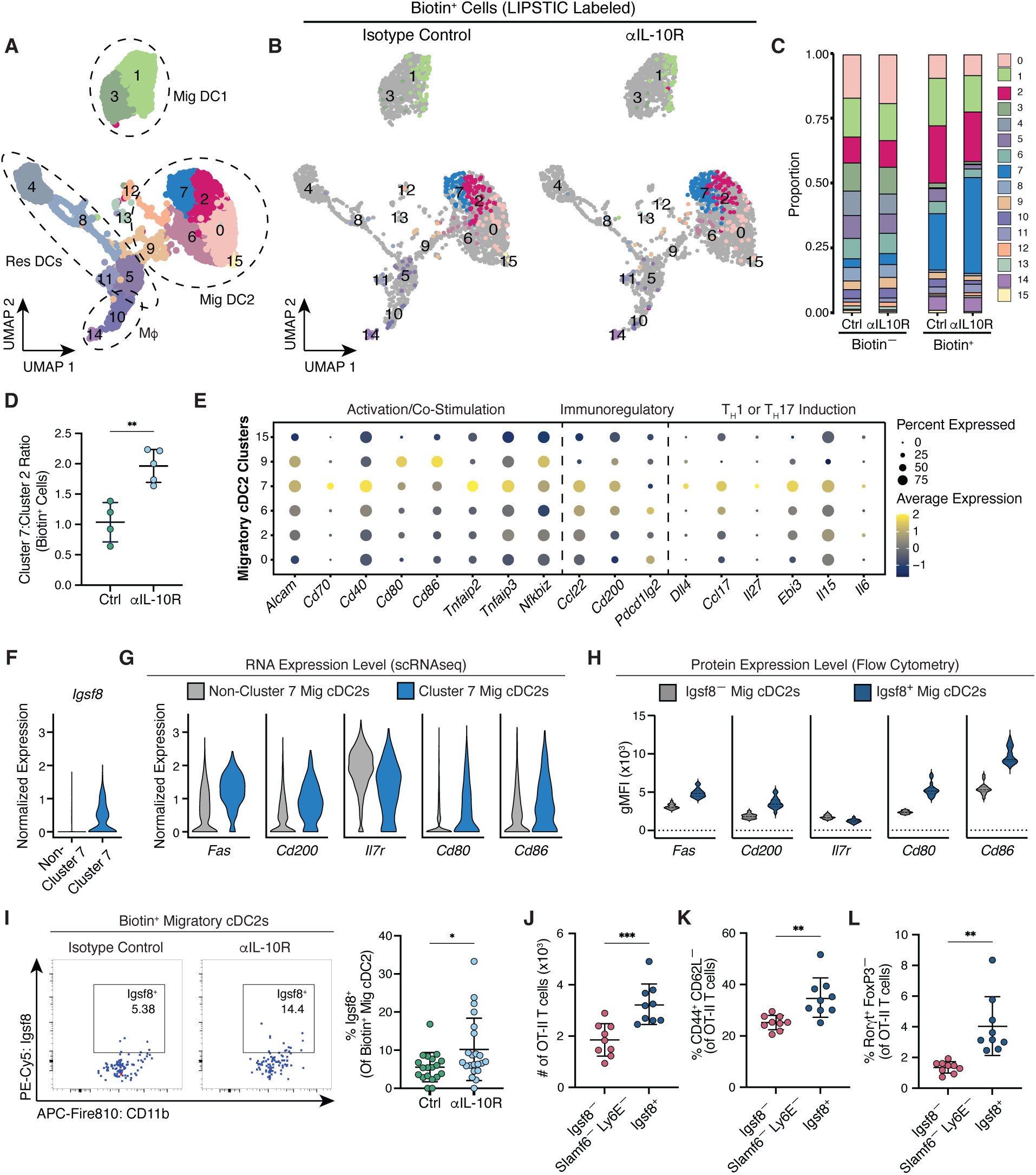
Inflammatory migratory cDC2s drive skewing of *A. muciniphila*-specific T cells towards alternative fates during inflammation. For (**A** to **G**), 1×10^6^ *Cd40lg*^SrtAv2^ Am124 T cells were adoptively transferred into ASF (untreated) and ASF+Akk mice 3 days after injection of 1 mg ɑIL-10R or isotype control antibodies. LIPSTIC labeling was performed 2.5 days later to sort and sequence *A. muciniphila*-presenting cells. (**A**) Uniform manifold approximation and projection (UMAP) plot of sorted and sequenced DCs from gLNs of one ASF mouse and control or ɑIL-10R-treated ASF+Akk mice (n = 4 to 5 mice per group). Cells are grouped into clusters via unsupervised hierarchical clustering. Dashed circles indicate annotation of cells based on expression of key marker genes as shown in fig. S8. (**B**) UMAP of sorted DCs separated by treatment group. Biotin^−^ cells are shown in gray. Biotin^+^ cells are colored according to which cluster they belong in. Biotin^+^ cells from ɑIL-10R-treated mice are downsampled to match the number of biotin^+^ cells from control animals. (**C**) Proportion of cells in each cluster among biotin^−^ and biotin^+^ cells. (**D**) The ratio of biotin^+^ cells in cluster 7 and biotin^+^ cells in cluster 2 between control and ɑIL-10R-treated mice. (**E**) Dot plot showing expression of genes differentially expressed among migratory cDC2 clusters. Size of dot indicates fraction of cells in each cluster that express gene; color of data indicates relative expression of gene. (**F**) Expression of *Igsf8* at the RNA level in non-cluster 7 and cluster 7 migratory cDC2s. (**G**) Expression of indicated genes at the RNA level in non-cluster 7 and cluster 7 migratory cDC2s. (**H**) gMFIs of indicated proteins in bulk Igsf8^−^ and Igsf8^+^ migratory cDC2s. Solid and dashed lines indicate median and quartiles (n = 4 to 6 mice per group, data is pooled from ASF and control or ɑIL-10-R-treated ASF+Akk mice, representative of four independent experiments). (**I**) 1×10^6^ *Cd40lg*^SrtAv2^ Am124 T cells were adoptively transferred into ASF (untreated) and ASF+Akk mice 3 days after injection of ɑIL-10R or isotype control antibodies. LIPSTIC labeling was performed 2.5 days later. Representative flow plots (left) and frequency (right) of Igsf8^+^ cells among biotin^+^ migratory cDC2s in control and ɑIL-10R-treated mice (n = 5 to 6 mice per group, per experiment, data pooled from four independent experiments). For (**J** to **L),** naive, CTV-labeled OT-II CD4^+^ T cells were co-cultured in the presence of exogenous OT-II peptide for 96 hours with Igsf8^−^ Slamf6^−^ Ly6E^−^ or Igsf8^+^ migratory cDC2 populations sorted from the gLNs of untreated ASF+Akk mice. **(J)** Total number of OT-II T cells after co-culture. **(K)** Frequency of activated (CD44^+^ CD62L^−^) OT-II T cells after co-culture. (**L**) Frequency of Rorγt^+^ FoxP3^−^ OT-II cells after co-culture. For (J to L), n = 4 to 5 mice per experiment, data pooled from two independent experiments. For (D), (I), and (J to L), each symbol represents one mouse and error bars represent mean and standard deviation. P-values were calculated by unpaired T-test for (D) and (I), and paired T-test for (J to L). Statistical significance denoted as *P < 0.05, **P < 0.01, ***P < 0.001, ****P < 0.0001.

Biotin^+^ migratory cDC2s occupied the same transcriptional clusters (clusters 2 and 7), both at steady state and during antibody blockade, but their distribution between the clusters shifted markedly. While at steady state the biotin^+^ cells were roughly evenly divided between the two clusters, during inflammation we observed a shift toward presentation by cells in cluster 7 (Fig. 5, C and D). Compared to cells in cluster 2, migratory cDC2s in cluster 7 were characterized by higher expression of co-stimulatory molecules (e.g., *Cd80, Cd86*, *Cd40* and *Cd70*) as well as genes previously implicated in T_H_1 and T_H_17 differentiation (e.g., *Dll4* and *Ccl17)* (*52–61*) (Fig. 5E), suggesting that they represent a more pro-inflammatory migratory cDC2 population.

To validate our scRNA-Seq results that *A. muciniphila* is more frequently presented by cluster 7 migratory cDC2s during inflammation, we used sc2markers (*62*) to identify distinguishing markers for cluster 7 migratory cDC2s that would enable detection of these cells by flow cytometry. Igsf8 was identified as the top marker for distinguishing cluster 7 from other migratory cDC2s, and we confirmed that its expression is restricted to cluster 7 migratory cDC2s (Fig. 5F, and Table S4). Flow cytometry analysis of Igsf8^+^ and Igsf8^−^ migratory cDC2s revealed that Igsf8^+^ cDC2s express many of the same markers as cluster 7 cDC2s indicating that they represent similar populations of cells (Fig. 5, G and H). Additionally, upon LIPSTIC labeling, a larger fraction of biotin^+^ migratory cDC2s was Igsf8^+^ in ɑIL-10R-treated animals than in the controls (Fig. 5I), consistent with the greater proportion of biotin^+^ cells in cluster 7 during IL-10R blockade.

Finally, to functionally test whether cluster 7 migratory cDC2s are more stimulatory and pro-inflammatory than other migratory cDC2s, we co-cultured naive ovalbumin-specific CD4^+^ OT-II T cells with Igsf8^+^ and Igsf8^−^ migratory cDC2s presenting ovalbumin peptide without exogenous T_H_-skewing cytokines. OT-II T cells cultured with Igsf8^+^ cDC2s upregulated Rorγt to a greater extent than those cultured with Igsf8^−^ cDC2s (fig. S9, A and B), suggesting that Igsf8^+^ cDC2s have an intrinsic ability to drive T_H_17 differentiation. However, we did not observe substantial differences in the total number, proliferation, and activation of OT-II T cells stimulated with Igsf8^−^ versus Igsf8^+^ cDC2s (fig. S9, C to E). We were concerned that non-cluster 2 cDCs could be confounding these comparative experiments, as Igsf8^−^ migratory cDC2s are heterogeneous and include cluster 9 cDCs, which were not labeled with biotin in our LIPSTIC experiments but express high levels of CD80 and CD86 (Fig. 5E). Therefore, to compare cluster 2 and cluster 7 migratory cDC2s more directly, we removed cluster 9 DCs from Igsf8^−^ migratory cDC2s based on their expression of Slamf6 and Ly6E (fig. S9, F and G, and Table S5). When compared to cluster 2-enriched Igsf8^−^ Slamf6^−^ Ly6E^−^ cDC2s, cluster 7-enriched Igsf8^+^ cDC2s induced increased numbers, proliferation, and activation of co-cultured OT-II T cells (Fig. 5, J and K, and fig. S9, H and I). Importantly, Igsf8^+^ cells maintained their superior ability to promote T_H_17 differentiation (Fig. 5L, and fig. S9, J and K). Altogether, these results show that during inflammation, increased presentation of commensal-derived antigens by pro-inflammatory DCs already existing at steady state can skew commensal-specific T cells towards alternative differentiation fates.

## Discussion

In this study, we show that migratory cDC2s prime *A. muciniphila*-specific T cells both at homeostasis, when the response against *A. muciniphila* is largely comprised of T_FH_ cells, and during inflammation, when the response also includes T_H_1 and T_H_17 cells. Several studies have documented heterogeneity amongst cDCs populations, especially within cDC2s, but how this diversity translates to functional consequences for the differentiation of T cells has been unclear (*26, 63–67*). Consistent with these previous reports, we find that cDC2s in the gLNs are heterogeneous and consist of multiple transcriptionally distinct subpopulations. By combining the LIPSTIC system with single-cell profiling and functional assays, we disentangled the contributions of specific cDC2 subpopulations in the priming of *A. muciniphila*-specific T cell responses. Overall, our results offer novel mechanistic insights into the basis of distinct adaptive immune responses to members of the microbiota.

At steady state, we find that *A. muciniphila*-specific T cells are primed by only some cDC2 populations, with the non-inflammatory cluster 2 and the pro-inflammatory cluster 7 cells contributing the most to antigen presentation. Surprisingly, the identity of the cDC2 populations that prime *A. muciniphila*-specific T cells during inflammation does not change; however, their relative proportions do change, with cluster 2 cells contracting and cluster 7 cells expanding, demonstrating how subtle shifts in the proportion of cDC2 subpopulations can dramatically affect T cell differentiation. Our results align with recent work showing that alterations in the composition of food-presenting APCs during helminth infection can disrupt food-specific T_REG_ induction (*36*).

Taken together, our findings support a model in which microbiota-reactive CD4^+^ T cells are primed by a heterogeneous pool of APCs and adopt particular differentiation trajectories through the integration of signals contributed by each subpopulation. This framework likely extends beyond the priming of microbiota-specific T cells, addressing the longstanding question of what dictates the nature of the CD4^+^ T cell response to a particular antigen. Our work also highlights the power of using LIPSTIC to track *in vivo* interactions. Detection of T cell interactions with multiple DC subpopulations would have been impossible with other approaches, as would capturing small changes in interactions across these subpopulations, as we have done here.

Given that a single cell type, cDC2s, can prime multiple diverse CD4^+^ T cell subsets (*22, 24, 42, 44, 68–73*), it is unlikely that each CD4^+^ T cell subset is instructed by an ontogenetically distinct type of APC (e.g., only cDC1s or cDC2s). As an alternative, it has been proposed that specialized subpopulations of cDC2s may each drive the differentiation of a distinct CD4^+^ subset, but this model has been challenging to assess due to limited tools to isolate these different cDC2 populations for *in vitro* assays or to selectively deplete them *in vivo*. Our data argue against a strict division of labor and instead suggest that a common set of cDC2 subpopulations, each with distinct functional properties, instructs the differentiation of multiple CD4^+^ T cell subsets, with the relative proportion of each subpopulation determining the differentiation trajectory. Crucially, we validate this model by isolating distinct cDC2 subpopulations and demonstrating their unique functional capacity *in vitro*. The development of more precise genetic tools will be important for defining the functional importance of different cDC2 subpopulations *in vivo*. It will also be essential to examine how widely applicable our model is by performing similar analyses for additional antigens in the gut as well as other anatomical sites.

In our dataset, the same cDC2 clusters were present in both untreated and αIL-10R-treated animals, indicating that large perturbations, such as inflammation, do not expand the range of possible cDC phenotypes. This observation aligns with previous studies showing that infection and cancer similarly fail to generate new cDC subpopulations(*36, 37*). Collectively, these data suggest that the programs or cues that drive cDCs towards all possible subpopulations may already be present in mice at homeostasis. This, in turn, raises the important question of what directs cDC2s toward a specific transcriptional identity and how perturbations, such as αIL-10R-induced inflammation, influence this process.

The prevailing model for cDC2 heterogeneity suggests that cDC2 phenotype is primarily shaped by environmental factors, with pre-cDC2s acquiring diverse phenotypic and functional traits depending on their tissue environment and microbial encounters. In line with this, the shift we observe during inflammation in the proportion of *A. muciniphila*-presenting cells in each transcriptional state may reflect differences in the local environment where these cells are sampling antigen and becoming activated. Supporting this idea, studies have shown that signals from the SI environment can modulate the phenotype and function of cDCs prior to their migration into the gLNs (*36, 74, 75*). At steady state, the integration of environmental signals, such as IL-10, along with microbe-derived signals causes cDC2s that interact with *A. muciniphila* to adopt the non-inflammatory cluster 2 phenotype. The role of IL-10 in skewing cDC2s towards an anti-inflammatory state is thus analogous to its role in enforcing a tolerogenic intestinal macrophage phenotype (*76, 77*). In the absence of IL-10 signaling (e.g., during IL-10R blockade), *A. muciniphila*-derived signals may instead drive cDC2s towards the pro-inflammatory cluster 7 phenotype. However, our results indicate that not all environmental signals can alter cDC polarization. The presence of SFB and the associated T_H_17-polarizing signals did not skew the differentiation of *A. muciniphila*-specific T cells, while blocking IL-10 signaling did. This difference suggests that environmental signals must meet certain requirements (e.g. reach a quantitative threshold or be of specific quality) to alter cDC phenotype.

An alternative model for cDC2 heterogeneity suggests that cDC2 subpopulations may represent ontogenetically distinct lineages. A recent study proposed that splenic cDC2A and cDC2B subsets, defined primarily by their expression of *Tbet*, *Esam*, and *Clec12a,* are ontogenetically determined lineages (*25*). The migratory cDC2s in our dataset express little to no *Tbet* or *Clec12a*, and while *Esam* expression was detected, it did not clearly segregate among migratory cDC2 clusters. Thus, whether the subpopulations that we observe in our dataset align with these previously described cDC2 lineages will require further investigation. Nonetheless, if the transcriptional clusters represent distinct lineages, how can the shift in antigen presentation across subpopulations be explained? One possibility is that inflammation alters the ability of different cDC subpopulations to acquire antigen, perhaps due to the differential localization of these subpopulations, or of *A. muciniphila* itself, within the intestines (*78*). Importantly, these two models of cDC heterogeneity are not mutually exclusive. In fact, it seems likely that both development programs and environmental signals contribute to the functional diversity of cDCs.

Our results provide mechanistic insight into our previous finding that *A. muciniphila*-specific T cell responses differ between gnotobiotic and conventionally-housed SPF animals (*4*). The increase in *A. muciniphila*-specific T_H_1 and T_H_17 cells that we previously reported in SPF mice mirrors the aIL-10R-induced changes observed here, suggesting that the varied T cell phenotypes observed in SPF mice may have been due to greater intestinal inflammation, and consequently increased antigen presentation by inflammatory cDC2s, compared to gnotobiotic animals. More broadly, our observations provide insight into the plasticity of commensal-specific T cell responses and how they can be shaped by inflammation. T cell responses against SFB are strongly biased towards a T_H_17 phenotype, even in the presence of a strong T_H_1-inducing pathogen, *Listeria monocytogenes* (*34*). However, responses against multiple other members of the microbiota, including *Clostridium* spp. and *H. hepaticus* appear to be malleable, with intestinal inflammation promoting differentiation of T_H_1 and/or T_H_17 cells against microbes that elicit regulatory responses at steady state (*5, 33, 79, 80*). Similar plasticity has been observed in the skin, where commensal-specific T cells shift their phenotype during tissue damage and inflammation (*81*). Altogether, these results suggest that T cell responses against most commensals may be responsive to contextual signals from the environment. Whether such plasticity can be broadly attributed to shifts in antigen presentation among heterogeneous subpopulations of APCs, as we have shown here, will need to be addressed in future studies.

Overall, this study advances our understanding of how microbiota-reactive T cells are primed and instructed towards distinct T_H_ fates by APCs. Our findings emphasize the critical importance of presentation by the appropriate proportions of phenotypically distinct APC populations to support homeostatic immune responses to the microbiota and demonstrate how inflammation can disrupt this balance, leading to aberrant, pro-inflammatory immune responses against the microbiota.

## Supporting information

Table S1 - Antibody List

Table S2 - Primer List

## Acknowledgements

We thank Russell Vance, Terri Laufer, and members of the Barton and Vance labs for comments on the manuscript and helpful discussions. We thank Hector Nolla, Alma Valeros, Kartoosh Heydari, Melaine Delcroix, and Harman Dhaliwal at the UC Berkeley Cancer Research Laboratory Flow Cytometry Facility for assistance with cell sorting and maintenance of flow cytometers. We thank the UC Berkeley Functional Genomics Laboratory for assistance with sequencing (RRID:SCR_022170). We thank Kaitlyn Siu for assisting in the maintenance of all gnotobiotic mouse lines and Keika Yan for technical assistance.

## Author contributions

Conceptualization: SLC, AL, GMB Formal analysis: SLC, AL

Funding acquisition: GMB Investigation: SLC, AL, AKL Methodology: SLC, AL, MCCC Resources: GDV, DM, GMB Supervision: GDV, DM, GMB Visualization: SLC, AL

Writing -- original draft: SLC, AL, GMB

Writing -- review & editing: SLC, AL, AKL, MCCC, GDV, DM, GMB

## Data and materials availability

Sequencing results will be deposited to the Gene Expression Omnibus public database (Note: at time of submission, the GEO database was not accepting new submissions, presumably due to the government shutdown. Once the database resumes normal operations, all sequencing results will be deposited). All other data are available in the main text or the supplementary materials. All materials (cell lines, mice, plasmids) are available upon request and after completion of a material transfer request to the corresponding author (GMB).

This article is subject to HHMI’s Immediate Access to Research policy, which requires that this article be made publicly available as initial and revised preprints deposited on a designated preprint server under a CC BY 4.0 license.

## Materials and Methods

### Animals

Mice were housed at the University of California, Berkeley under specific-pathogen-free (SPF) or gnotobiotic conditions. Male and female mice between 6-12-weeks of age were used for experiments, except in the case of bone marrow chimera experiments in which mice were analyzed 8-12 weeks after bone marrow transplant. All experiments were performed in accordance with the Animal Care and Use Committee guidelines at the University of California, Berkeley. Amuc124 TCR transgenic mice were generated and maintained in our laboratory (*4*). *Rag1^-/-^* (strain number 002216), Thy1.1 (strain number 000406), CD45.1 (Strain number 002014), *Ccr7^-/-^* (strain number 006621), JAXBoy (strain number 033076), and 7B8 TCR transgenic (strain number 027230) mice were purchased from Jackson laboratories and maintained in our facility. Δ1+2+3 (cDC2 deficient) mice were provided by Kenneth Murphy (*39*). *Zbtb46*-DTR (strain number 019506) and *Batf3^-/-^* (cDC1 deficient; strain number 013755) mice were provided by Michel DuPage. *H2-Ab1^-/-^* (Taconic Model No. ABBN12) and OT-II TCR transgenic mice (Taconic Model No. 11490) were provided by Ellen Robey. *Cd40*^G5/G5^ and *Cd40lg*^SrtAv2^ were provided by Gabriel Victora (*27*).

Amuc124 TCR transgenic mice were bred to *Rag1^-/-^* and Thy1.1 mice to generate Amuc124 *Rag1^-/-^* Thy1.1 mice. 7B8 TCR transgenic mice were bred to *Rag1^-/-^*and CD45.1 mice to generate 7B8 *Rag1^-/-^* CD45.1 mice. *Cd40lg*^SrtAv2^ were bred to Amuc124 *Rag1^-/-^* Thy1.1 mice to generate Amuc124 *Cd40lg*^SrtAv2^ Rag1^-/-^ Thy1.1 mice. All Amuc124 TCR Tg lines were maintained free of *A. muciniphila*, as confirmed by PCR (described below). *Cd40*^G5/G5^ mice were crossed to *H2-Ab1^-/-^* mice to generate *Cd40*^G5/G5^*H2-Ab1^-/-^* mice. *Cd40*^G5/G5^ mice were re-derived to germ-free status by cesarean section and pups were fostered onto germ-free Swiss Webs.

Gnotobiotic C57BL/6NTac mice colonized with altered Schaedler’s flora (ASF) were obtained from Taconic Biosciences and imported into the gnotobiotic facility at the University of California, Berkeley. ASF+Akk mice were previously generated from C57BL/6NTac ASF mice by two oral gavages of 10^9^ CFUs of *A. muciniphila* (Berkeley strain previously isolated from our SPF colony (*4*)) two days apart. ASF+Akk+SFB mice were generated from germ-free C57BL/6NTac mice by two oral gavages with fecal supernatant from mice monocolonized with SFB (frozen feces acquired from Dan Littman). SFB-monocolonized mice were then co-housed with ASF+Akk mice to generate ASF+Akk+SFB mice. OMM^12^ mice were generated by gavaging germ-free C57BL/6NTac once with OMM^12^ cecal supernatant (frozen cecal contents provided by Gabriel Victora). OMM^11^ mice were generated by treating OMM^12^ mice with a six-week course of colistin sulfate in the drinking water (2 g/L colistin sulfate, 3% sucrose, Cayman Chemical). *A. muciniphila* and SFB colonization was detected by 16S qPCR on fecal DNA extracts, while OMM^12^ microbiota colonization was detected by endpoint PCR on fecal DNA extracts, as described below. Absence of bacterial contaminants was routinely tested by bacterial plating and fecal V3-V4 16S rDNA sequencing (Zymo Research). ASF, ASF+Akk, ASF+Akk+SFB and OMM^12^ mouse colonies were maintained in separate flexible film isolators (Class Biologically Clean). OMM^11^ mice were maintained in sterilized Isocages on the Techniplast Isocage P rack system. For experiments with gnotobiotic mice, animals were removed from gnotobiotic isolators and transferred into sterilized Isocages (Tecniplast) within a biosafety cabinet. Cages were then housed on a Tecniplast Isocage P rack system. In some experiments with ASF mice, we detected contamination with *Blautia coccoides* and *Enterocloster clostridioformis* (members of the OMM^12^ microbiota). We have confirmed that the presence of these two bacterial species does not alter the differentiation of *A. muciniphila*- specific T cells (fig. S10).

C57BL/6J SPF *A. muciniphila*-negative mice (referred to as C57BL/6J SPF mice unless otherwise noted) were generated by treating mice with a three week course of doxycycline hyclate in the drinking water (3 g/L doxycycline hyclate, 1% sucrose, pH = 7, Sigma). C57BL/6J SPF+Akk mice were generated from C57BL/6J SPF mice by a single oral gavage of 10^9^ CFUs of *A. muciniphila* (Berkeley strain). Consistent with prior work, we observed variable *A. muciniphila*-specific T cell responses in our C57BL/6J SPF+Akk mice. We therefore performed experiments using mice from a line established from C57BL/6J SPF+Akk mice screened for robust *A. muciniphila*-specific T cell responses.

ASF and ASF+Akk *Cd40*^G5/G5^ mice were generated by cohousing with C57BL/6NTac ASF and ASF+Akk mice. OMM^11^ and OMM^12^ *Cd40*^G5/G5^ were generated by cohousing germ-free *Cd40*^G5/G5^ with C57BL/6NTac OMM^11^ and C57BL/6NTac OMM^12^ mice. SPF *Cd40*^G5/G5^ were generated by cohousing germ-free *Cd40*^G5/G5^ with C57BL/6J SPF mice. SPF+Akk *Cd40*^G5/G5^ were generated by cohousing germ-free *Cd40*^G5/G5^ with C57BL/6J SPF mice followed by two oral gavages of 10^9^ CFUs of *A. muciniphila* (Berkeley strain). All experiments were performed on mice that acquired their microbiota from birth via vertical colonization, except for in fig. S10 in which mice were gavaged with *B. coccoides* and *E. clostridioformis*.

### Tissue processing for flow cytometry

For isolation of T cells from the gut-draining lymph nodes and spleen, cells were isolated by mechanical dissociation on a 70-µm filter. For the spleen, cells were treated with ACK lysis buffer to remove red blood cells prior to staining for flow cytometry. For isolation of T cells from Peyer’s patches, cells were isolated by mechanical dissociation on a 100 mm filter. For isolation of dendritic cells (or T cells for when Am124 T cells were examined less than 7 days post-transfer) from gut-draining lymph nodes, tissue was cut into small pieces and digested in RPMI containing 2% (v/v) FBS, 20 mM HEPES (Gibco), and 400 U/mL collagenase IV (Worthington) for 25 minutes at 37°C prior to mechanical dissociation on a 70-µm filter. In experiments where Thetis cells and innate lymphoid cells were examined in addition to dendritic cells (Fig. 3, C to E, and fig. S3. A to C, and F to H), gut-draining lymph nodes were digested in RPMI containing 5% (v/v) FBS, 100 U/mL penicillin (Gibco), 100 µg/mL streptomycin (Gibco), 1X Glutamax (Gibco), 10 mM HEPES, 1 mg/mL collagenase A (Roche), and 1 U/mL DNaseI (Sigma) instead of collagenase IV.

For isolation of cells from small intestine lamina propria (after removal of Peyer’s patches) and large intestine lamina propria (cecum and colon, after removal of cecal patch), the intestinal tissue was flushed with PBS, cut longitudinally, washed in PBS to remove mucus and intestinal contents, then cut into small pieces. The small intestines were then incubated at 37°C for 20 min with stirring in Hank’s balanced salt solution (HBSS) with 1 mM DTT, 10% (v/v) FBS, 100 U/mL penicillin, 100 µg/mL streptomycin, 1X Glutamax and 10 mM HEPES. This was followed by incubating the tissue at 37°C for 25 min with stirring in HBSS with 1.3 mM EDTA, 100 U/mL penicillin, 100 µg/mL streptomycin, 1X Glutamax and 10 mM HEPES, with additional dissociation of epithelial cells by shaking in 10 mL PBS. For large intestines, DTT and EDTA dissociations were performed as one step in HBSS with 1 mM DTT, 1.3 mM EDTA, 100 U/mL penicillin, 100 µg/mL streptomycin, 1X Glutamax and 10 mM HEPES for 30 minutes. For both small and large intestines, tissues were digested in RPMI with 1 mg/mL collagenase VIII (Sigma), 5 µg/mL DNaseI, 100 U/mL penicillin, 100 µg/mL streptomycin, 1X

Glutamax and 10 mM HEPES for 45 min. Any tissue pieces remaining after the digestion were mechanically dissociated on a 100 µm filter. Finally, lymphocytes were collected at the interface of a 44%/67% Percoll gradient (GE Healthcare).

### Flow cytometry

Cells were isolated as described above and stained in a 96-well U-bottom plate. Dead cells were excluded with a fixable live/dead dye in PBS. Prior to surface staining, cells were blocked with 2.5 µg/mL Fc block (anti-CD16/32). Surface staining was performed in PBS with 2% FBS (v/v), 2 mM EDTA and 0.1% NaN_3_ (excluded when sorting for live cells or staining for CXCR5) for 25 minutes at 4° C, or for 1 h at room temperature (RT) when staining for CXCR5. Intracellular transcription factor staining (eBioscience Foxp3/Transcription Factor Staining Buffer Set, Invitrogen) was performed by fixation for 40 minutes at RT followed by staining for 40 min at RT. To obtain cell counts, 15 to 20 uL of CountBright Absolute Counting Beads (Invitrogen) were added to samples just prior to running on the cytometer. Samples were analyzed on a BD LSR Fortessa X-20, BD Symphony A3 or Cytek Aurora and analyzed using the FlowJo Software package. The list of antibodies used can be found in Table S1.

### Igsf8 antibody conjugation for flow

The Igsf8 antibody (Biotechne) we used for flow cytometry was conjugated to a PE-Cy5 fluorophore using the Lightning Link Conjugation Kit (Abcam) according to the manufacturer’s protocol.

### Adoptive T cell transfers

Inguinal and gut-draining lymph nodes and spleen were harvested from TCR transgenic mice (Am124 or 7B8) and dissociated mechanically on a 70 µm filter. T cells were isolated by negative selection using the StemCell EasySep Mouse T Cell isolation kit (STEMCELL Technologies) according to the manufacturer’s protocol. The purity of T cells was verified by flow cytometry (Live TCRβ^+^ CD4^+^ CD8^−^ CD62L^+^ CD44^−^). Purity was generally 85-95%. T cells were transferred via retroorbital injection into recipient mice with isoflurane anesthesia. The number of cells transferred in each experiment is indicated in the figure legend. For experiments with low frequency transfers (1 x 10^4^ cells per mouse), naïve Am124 CD4 T cells (Live TCRβ^+^ CD4^+^ CD8^−^ CD62L^+^ CD44^−^) were sorted on a BD FACSAria Fusion or a Cytek Aurora CS cell sorter.

### SrtA substrate

Biotin–aminohexanoic acid–LPETGS (C-terminal amide, 95% purity) was purchased from LifeTein (custom synthesis). Stock solutions were prepared at 20 mM in sterile PBS.

### LIPSTIC *in vivo*-labeling experiments

1×10^6^ Amuc124 *Cd40lg*^SrtAv2^ *Rag1^-/-^* Thy1.1 T cells were transferred into *Cd40*^G5/G5^ mice via retroorbital injection. 64 hours later, 2 μM (100 μl of 20 mM solution) Biotin–LPETG substrate was injected into *Cd40*^G5/G5^ mice intraperitoneally vsix times over two hours (20 minute intervals between injections). In total, each mouse received 12 μM of substrate. Control mice that did not receive LPETG substrate were injected with PBS at the same intervals. Gut-draining lymph nodes were collected 40 min after the last injection.

### Bone marrow chimeras

Mice were treated intraperitoneally with 75 mg/kg busulfan (Cayman Chemical) provided as daily doses of 25 mg/kg for three consecutive days. Two days after the last dose of busulfan, 10 x 10^6^ donor bone marrow cells from femurs and tibiae were transferred via retroorbital injection. Mice were analyzed 8-12 weeks after bone marrow transplant. In each bone marrow transfer experiment, one mouse was reconstituted with JAXBoy (CD45.1) bone marrow instead of C57BL/6J bone marrow to verify efficient replacement of the hematopoietic compartment.

These mice are included in figures along with mice reconstituted with C57BL/6J bone marrow as a wild-type control. For bone marrow chimera experiments in Fig. 3, F and G and fig. S4, B and F, ASF control mice were not bone marrow chimeras. In one of two experiments in fig. S2E, ASF control mice were not bone marrow chimeras. For experiments in fig. S4, A and B, bone marrow chimeras were treated daily with 20 ng/g diphtheria toxin (Sigma) starting two days prior to transfer and until endpoint analysis at 1.5 days post-transfer.

### Colonization with bacterial strains

*A. muciniphila* (Berkeley strain previously isolated from our SPF colony) (4) was grown in liquid mucin medium containing 0.4 g/L KH_2_PO_4_, 0.53 g/L Na_2_HPO_4_, 0.3 g/L NH_4_Cl, 0.3 g/L NaCl, 0.1 g/L MgCL_2_·6H2O, 0.15 g/L CaCl_2_·2H_2_O, 4 g/L NaHCO_3_, 0.45 g/L L-Cys·HCl·H_2_O, 2.5 g/L type III hog gastric mucin (Sigma), 3 mL/L trace mineral solution (ATCC) and 1 mg/L resazurin (Sigma). To prevent the growth of contaminants, the growth medium was supplemented with 10 µg/mL gentamicin and 12 µg/mL kanamycin. *B. coccoides* YL58 and *E. clostridioformis* YL32 were obtained from the Leibniz Institute DSMZ and grown in Anaerobic Medium containing 18.5 g/L brain heart infusion, 5 g/L yeast extract, 15 g/L trypticase soy broth, 2.5 g/L KH_2_PO_4_, 1 mg/L haemin, 0.5 g/L glucose, 0.4 g/L Na_2_CO_3_, 0.5 g/L L-Cys·HCl·H_2_O, 5 mg/L menadione, 3% heat-inactivated fetal bovine serum, and 1 mg/L resazurin. All bacterial strains were grown at 37° C in an anaerobic chamber (Coy Laboratory) containing 5% hydrogen, 5% carbon dioxide and 90% nitrogen until saturation. Bacteria were then harvested by centrifugation at 6,000 g at 4° C for 10 minutes and washed once with PBS before resuspension in anaerobic PBS at a concentration of 1×10^10^ CFUs/mL. Mice were intragastrically gavaged with 100 µL (1×10^9^ CFUs total) of the bacterial resuspension using a smooth ball tip feeding needle. For gavages with fecal (SFB) and cecal (OMM^12^) bacterial supernatants, frozen fecal and cecal contents were homogenized through a 100 µm filter with sterile PBS and centrifuged at 200 g at RT for 5 minutes to pellet large particles. Mice were intragastrically gavaged with 100 µL of the bacteria-containing supernatant using a smooth ball tip feeding needle.

### Detection of bacterial strains in feces

For the detection of *A. muciniphila*, SFB, and OMM^12^ bacteria, including *B. coccoides* and *E. clostridioformis*, DNA was isolated from fresh or frozen fecal pellets using the QIAamp Fast DNA Stool Minikit (Qiagen) or the DNeasy PowerSoil Pro QIAcube Kit (Qiagen) for *A. muciniphila*, the DNeasy PowerLyzer PowerSoil Kit (Qiagen) for SFB and OMM^12^ bacteria, or the DNeasy PowerSoil Pro QIAcube Kit (Qiagen) for *B. coccoides* and *E. clostridioformis*. DNA extractions were performed using a QIAcube Connect according to the manufacturer’s instructions. For the DNeasy PowerLyzer PowerSoil Kit, the following modifications were made: prior to homogenizing the fecal pellets, samples (feces with Bead Solution and C1 solution) were incubated for 10 minutes at 65° C followed by 10 minutes at 95° C. After adding C2 solution, samples were incubated on ice for 5 minutes prior to centrifugation. After adding C3, samples were incubated on ice for 10 minutes prior to centrifugation. After loading the DNA onto the spin column, DNA was washed with 100% ethanol prior to the two washes with C5 solution. Homogenization was performed on a PowerLyzer 24 Homogenizer (Qiagen). To quantify *A. muciniphila* and SFB genome amounts, a standard curve was generated using the entire 16S gene for *A. muciniphila* or a portion of the 16S gene for SFB cloned into a pCR-BluntII TOPO vector (Invitrogen). Quantitative PCR was performed on a StepOnePlus thermocycler (Applied Biosystems) with SsoAdvanced universal SYBR green supermix (Bio-RAD) or PowerUp SYBR Green Master Mix (Applied Biosystems) for *A. muciniphila* and FastStart Taqman Probe Master Mix (Sigma) for SFB. Endpoint PCR for the detection of OMM^12^ bacteria was performed on a C1000 Touch Thermal Cycler (Bio-Rad) with DreamTaq PCR Master Mix (Thermo). Primers used to detect OMM^12^ bacteria were modified versions of previously published primer sets (*37*). All primers and probes were generated by Integrated DNA Technologies. The list of primers can be found in Table S2.

### Anti-IL-10R treatment

Animals were injected intraperitoneally with 1 mg anti-mouse IL-10R antibody (BioXCell BE0050) or IgG1 isotype control (BioXCell BE0088) diluted in PBS. In some experimental replicates, control animals were injected with PBS alone rather than the isotype control antibody. Three days after antibody treatment, T cells were transferred via retroorbital injection (1 x 10^4^ Am124 cells for Am124 T cell differentiation; 2.5 x 10^5^ Am124 cells for Am124 activation in gLNs;1 x 10^6^Am124 *Cd40lg*^SrtAv2^ cells for LIPSTIC labeling). For T cell differentiation studies, antibody injections continued once per week for the duration of the experiment, and animals were sacrificed 12 days after the T cell transfer for tissue harvest. For LIPSTIC labeling experiments, labeling was performed 64 hours post T cell transfer (as described above).

### Segmentation of gut-draining lymph nodes

The gut-draining lymph nodes were identified as described previously (*29–31*). The duodenal lymph nodes included the hepatic-celiac lymph node located underneath the portal vein and near the liver, the pancreatic-duodenal lymph node embedded in the pancreas and the uppermost lymph node of the mesenteric lymph node chain (closest to the stomach). The jejunal lymph nodes included the next lower two to three lymph nodes after the duodenal lymph node in the mesenteric lymph node chain and the ileal lymph nodes included the next lower two to three lymph nodes after the jejunal lymph nodes. The cecal-colonic lymph nodes included the lowermost lymph node of the mesenteric lymph node chain (closest to the cecum) and a small ascending-colonic lymph node generally located in the mesentery between the ascending colon and the mesenteric lymph node chain.

### *In vitro* DC–T cell co-culture

CD4^+^ OT-II T cells were harvested from the spleen and gLNs of OT-II mice, labeled with 0.5 uM CellTrace Violet (Invitrogen) according to the manufacturer’s instructions and then stained with antibodies, as described above, for sorting. 750 naive, CTV-labeled CD4+ OT-II T cells (Live TCRβ^+^ CD4^+^ CD8^−^ CD62L^+^ CD44^−^) were sorted into U-bottom, TC-treated 96-well plates containing RPMI supplemented with 10% (v/v) FBS, 100 U/mL penicillin, 100 µg/mL streptomycin, 1X Glutamax, 10 mM HEPES, and 2% (v/v) of T-stim media (VWR). cDCs were harvested from pooled gLNs and stained with antibodies, as described above, for sorting. Following staining, cDCs were washed and resuspended in PBS with 10% FBS (v/v), 2 mM EDTA and 5 nM Sytox Green Nucleic Acid Stain (Thermo). 150 Igsf8^−^, Igsf8^−^ Slamf6^−^ Ly6E^−^ or Igsf8^+^ migratory cDC2s (Live TCRβ^−^ CD19^−^ CD64^−^ Ly6C^−^ MHCII^hi^ CD11c^int^ CD11b^+^ XCR1^−^) were sorted into wells with sorted naive, CTV-labeled CD4^+^ OT-II T cells. Following sorting of both T cells and cDCs, 1 µM of OT-II peptide (GenScript) was added to wells and plates were then spun down at 400 rpm for 3 minutes at RT to physically cluster the cells. Cells were incubated at 37° C, 5% CO_2_ for 96 hours before antibody staining, as described above. Fresh, congenically marked splenocytes (CD45.1^+^) were added to wells prior to staining to prevent cell loss. Sorting was performed on a Cytek Aurora CS cell sorter.

### Preparation of dendritic cells for scRNA sequencing

64 hours after transferring Amuc124 *Cd40lg*^SrtAv2^ *Rag1^-/-^* Thy1.1 T cells into ASF+Akk *Cd40*^G5/G5^ mice, *in vivo* LIPSTIC labeling was performed as described above. 40 minutes after the final Biotin–LPETG injection, gut-draining lymph nodes were harvested and a single cell suspension was prepared as described above (digestion with collagenase IV followed by mechanical dissociation on a 70 µm filter). Dendritic cells were enriched using the EasySep Mouse CD11c Positive Selection Kit II (STEMCELL Technologies) according to the manufacturer’s instructions. Samples were stained with the following combination of fluorescent anti-mouse antibodies for sorting and anti-mouse TotalSeq-B antibodies for 30 minutes at 4°C: APC-Cy7 anti-Thy1.2, APC-Cy7 anti-CD19, APC-Cy7 anti-B220, BV421 anti-MHCII, PE anti-Biotin, TotalSeq-B anti-XCR1, TotalSeq-B anti-CD11b. In a second step, cells were stained with TotalSeq-B anti-PE to enable detection of biotinylated LIPSTIC-labeled cells by sequencing.

Cells from each individual mouse were also labeled with a unique TotalSeq-B hashtag antibody to allow for multiplexing of samples from several mice on one 10X lane. Following staining, cells were washed and resuspended in PBS with 1% FBS (v/v), 2 mM EDTA, 0.1% NaN_3_ and 0.5 µM Sytox Green Nucleic Acid Stain (Thermo). Dendritic cells (Live Thy1.2^−^ B220^−^ CD19^−^ CD11c^+^ MHCII^+^) were sorted using a 130-µm nozzle on the BD FACSAria Fusion cell sorter. Biotin^+^ and biotin^−^ cells were sorted separately and then all biotin^+^ cells were combined with a portion of the Biotin^−^ cells from each mouse to enrich for biotin^+^ cells.

### Library preparation for single cell RNA sequencing

scRNA-seq libraries of sorted dendritic cells were generated on the 10X Genomics Single Cell Chromium System according to the manufacturer’s instructions. The dataset used in Figure 3 was generated using a Chromium Next GEM Single Cell 3’ Kit v3.1 along with the 3’ Feature Barcode Kit. The dataset used in Figure 4 was generated using a GEM-X Universal 3’ Gene Expression v4 kit along with the Chromium GEM-X Single Cell 3 Feature Barcode Kit v4. Both libraries were sequenced on the Illumina NovaSeq 6000 at the UC Berkeley Functional Genomics Laboratory.

### Single-cell RNA-Seq analysis

Raw fastq files were aligned to the Mouse (GRCm39) 2024-A genome with 10X Genomics Cell Ranger v9.0.0 using 10X Genomics Cloud Analysis with the “Include introns=True” option (*82*). The Cell Ranger aggregation output (filtered_feature_bc_matrix directory) was used as input for the Seurat package (Version 5.2.1) (*83–87*). Cells with a high percentage (>5%) of reads aligning to mitochondrial genes were excluded as likely dying cells.

Low quality cells with poor gene capture (<700 genes for the dataset in Figures 3 and S5, <2500 genes for the dataset in Figures 5 and S8) were also excluded. Doublets were identified as cells containing multiple hashtag antibodies and excluded. The dataset was normalized using the LogNormalize method and scaled using the ScaleData function in Seurat. Protein expression data from hashtag antibodies was normalized using the NormalizeData function in Seurat with CLR (centered log ratio transformation) as the normalization method. Linear dimensional reduction was performed on the top 2000 variable genes. The top 20-30 principal components were used for clustering using the standard Seurat workflow. For the steady state dataset, the Clustree algorithm (*88*) was used to generate cluster trees to determine the best resolution for clustering the data. Annotation of clusters was performed based on the expression of cell type specific markers as shown in figures S5 and S8. Differential expression was performed using the FindMarkers functions in Seurat with the Wilcoxon Rank Sum test, and P-values were adjusted with the Bonferroni correction for multiple hypothesis testing. Volcano plots for figure S8 were generated using TidyPlots (*89*). The surface markers for the migratory cDC2 clusters in Figure 9 and S10 were identified using the Detect_single_marker function in the sc2marker package (*61*).

For the correlation analysis in Figure 3K, cells were subsetted to include only migratory cDC2s. These were defined as cells in clusters 0, 1, 4, 5, and 8 that had normalized expression of Xcr1 protein (based on Hashtag antibody) below 0.5 and normalized expression of Ccr7 (RNA) above 1. The correlatePairs function in the scran package (*90*) was used to calculate the Spearman’s rank correlation between the normalized Biotin signal and scaled expression data for every gene. For the gene set enrichment analysis (GSEA) (*91*) on significantly correlated genes, the msigdbr package (R version 7.5.1) (*92*) was used to download Hallmark and Gene Ontology gene sets (*93, 94*) and the fgsea package (*95*) was used to run the GSEA in R. For fgsea, minSize was set at 10 and maxSize was set at 500.

For the inflammation dataset, there were several clusters of contaminating non-DCs that were removed prior to clustering analysis on DCs (and macrophages) alone. These included pDCs, NK cells, B cells, and T cells. pDCs were identified as cells expressing *SiglecH*, *Tcf4*, and *Ly6D*; NK cells were identified as expressing *Ncr1*, *Gzma*, *Klrb1c*, *Nkg7*; B cells were identified as expressing *Bank1*, *Igkc*, *Jchain*, *Ighm*, *Ighd*, *CD79a*, *CD79b*; T cells were identified as expressing *Trac*, *Trbc1*, *Trbc2*, *Skap1*.

Before generating the UMAP for figure 5B, the number of biotin^+^ from the mice treated with aIL-10R was reduced by randomly sampling until the number of cells matched the number of biotin^+^ cells from the isotype control treated animals.

### Quantification and statistical analysis

Statistical tests were performed as indicated in the figure legends with Prism 10 software (Graphpad Prism)

---

**Supplementary Figure 1:**
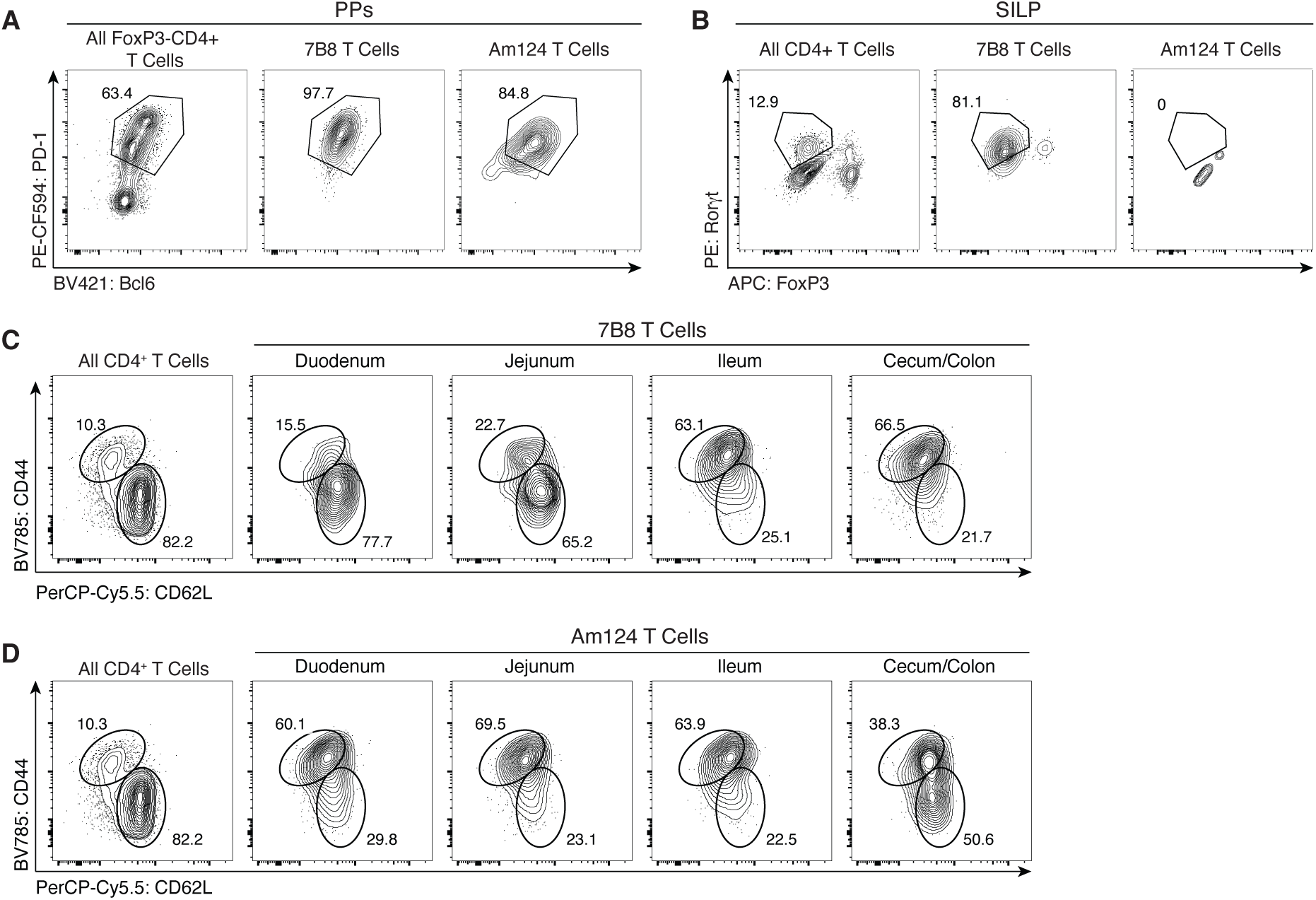
Expression of differentiation and activation markers by 7B8 and Am124 T cells. (**A**) Representative flow plots showing expression of T_FH_ markers (Bcl6 and PD- 1) by endogenous (left), 7B8 (center), and Am124 (right) T cells in the PPs of ASF+Akk+SFB mice. (**B**) Representative flow plots showing expression of T_H_17 markers (Rorγt^+^ FoxP3^−^) by endogenous (left), 7B8 (center), and Am124 (right) in the SILP of ASF+Akk+SFB mice. (**C**) Representative flow plots showing activated (CD44^+^ CD62L^−^) 7B8 T cells in segmented gLNs. (**D**) Representative flow plots showing activated (CD44^+^ CD62L^−^) Am124 T cells in segmented gLNs. **A**

**Supplementary Figure 2:**
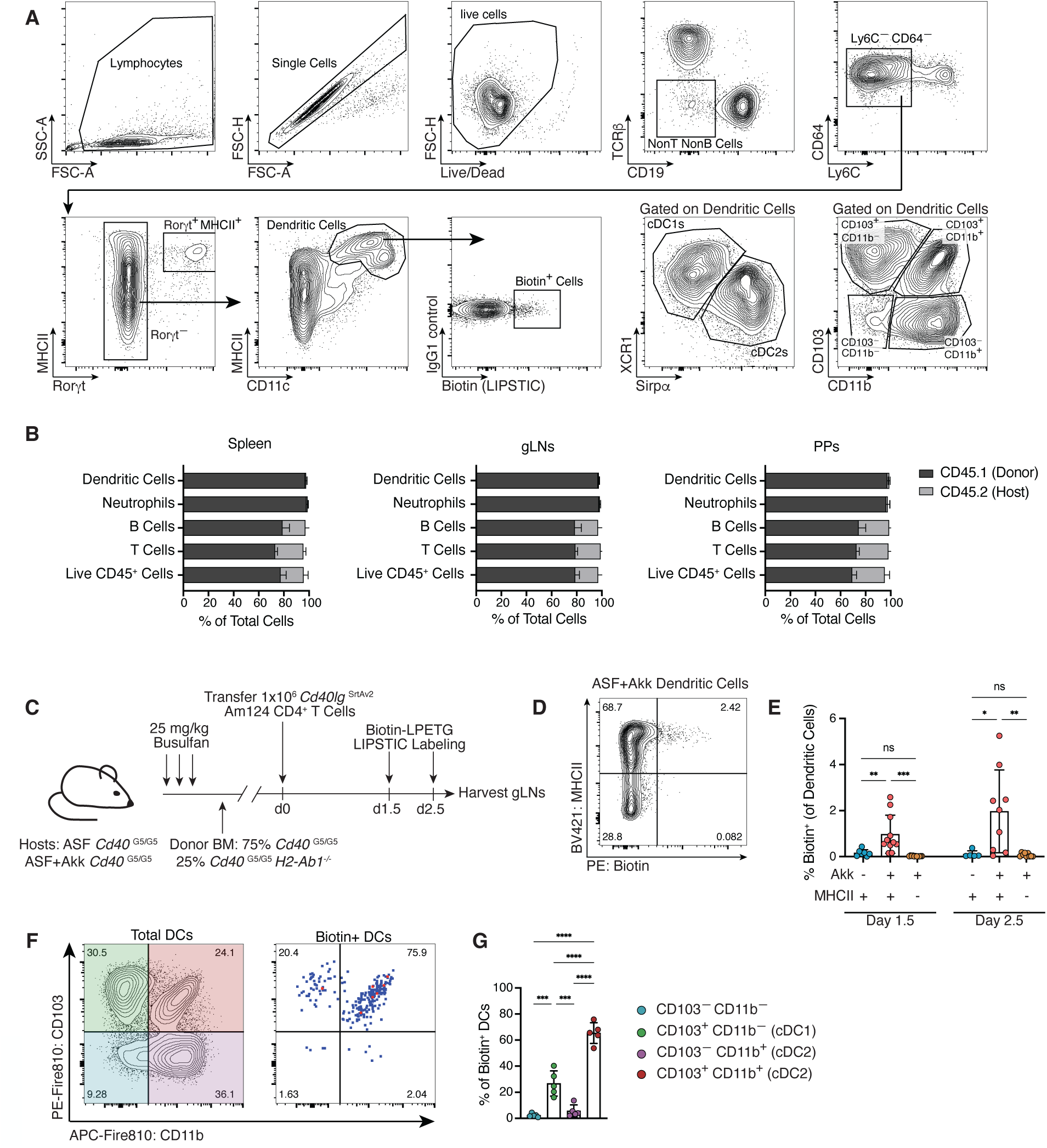
Additional characterization of the Am124 LIPSTIC-labeling system. (**A**) Flow plots showing the general gating scheme for cDCs. (**B**) C57BL/6N (CD45.2^+^) mice were treated with 25 mg/kg busulfan every day for three days (75 mg/kg total), followed by reconstitution with JAXBoy (CD45.1^+^) bone marrow. Spleen (left), gLNs (middle) and PPs (right) were harvested 8 to 12 weeks later. Plots show frequency of CD45.2^+^ (host) and CD45.1^+^ (donor) cells out of total live cells, T cells, B cells, neutrophils, and cDCs. Error bar represents mean and standard deviation (n = 5 mice per group, data is representative of two independent experiments). (**C**) Experimental timeline for (**D** and **E**). BMCs were generated by treating mice with 25 mg/kg busulfan daily for three days (75 mg/kg total), followed by reconstitution with 75% *Cd40*^G5/G5^ bone marrow and 25% *Cd40*^G5/G5^ *H2-Ab1*^-/-^ bone marrow. After 8 to 12 weeks, 1x 10^6^ *Cd40lg*^SrtAv2^ Am124 T cells were adoptively transferred and LIPSTIC labeling was performed 1.5 and 2.5 days post transfer. (**D**) Representative flow plot showing LIPSTIC labeling of MHCII^+^ and MHCII^−^ cDCs in mixed BMCs. cDCs were gated as Live Thy1.2^−^ F4/80^−^ SiglecF^−^ Ly6G^−^ CD19^−^ CD64^−^ CD11c^+^ to avoid gating with MHCII. (**E**) Frequency of labeled biotin^+^ cDCs in the gLNs of ASF and ASF+Akk mixed BMCs at 1.5 and 2.5 days post transfer. For ASF+Akk mice, cells are divided based on the expression of MHCII. All mice received LPETG substrate. For (C to E), n = 3 to 7 mice per group, per experiment; data is pooled from two independent experiments. (**F**) Representative flow plots showing expression of CD11b and CD103 on total (left) and biotin^+^ cDCs (right) in gLNs. (**G**) Percentage of biotin^+^ DCs expressing CD103 and CD11b. For (F to G), n = 3 to 5 mice per group; data is representative of three independent experiments. For all plots, each symbol represents one mouse and error bars represent mean and standard deviation. P-values were calculated using one-way ANOVA. Statistical significance denoted as not significant (ns), *P < 0.05, **P < 0.01, ***P < 0.001, ****P < 0.0001.

**Supplementary Figure 3:**
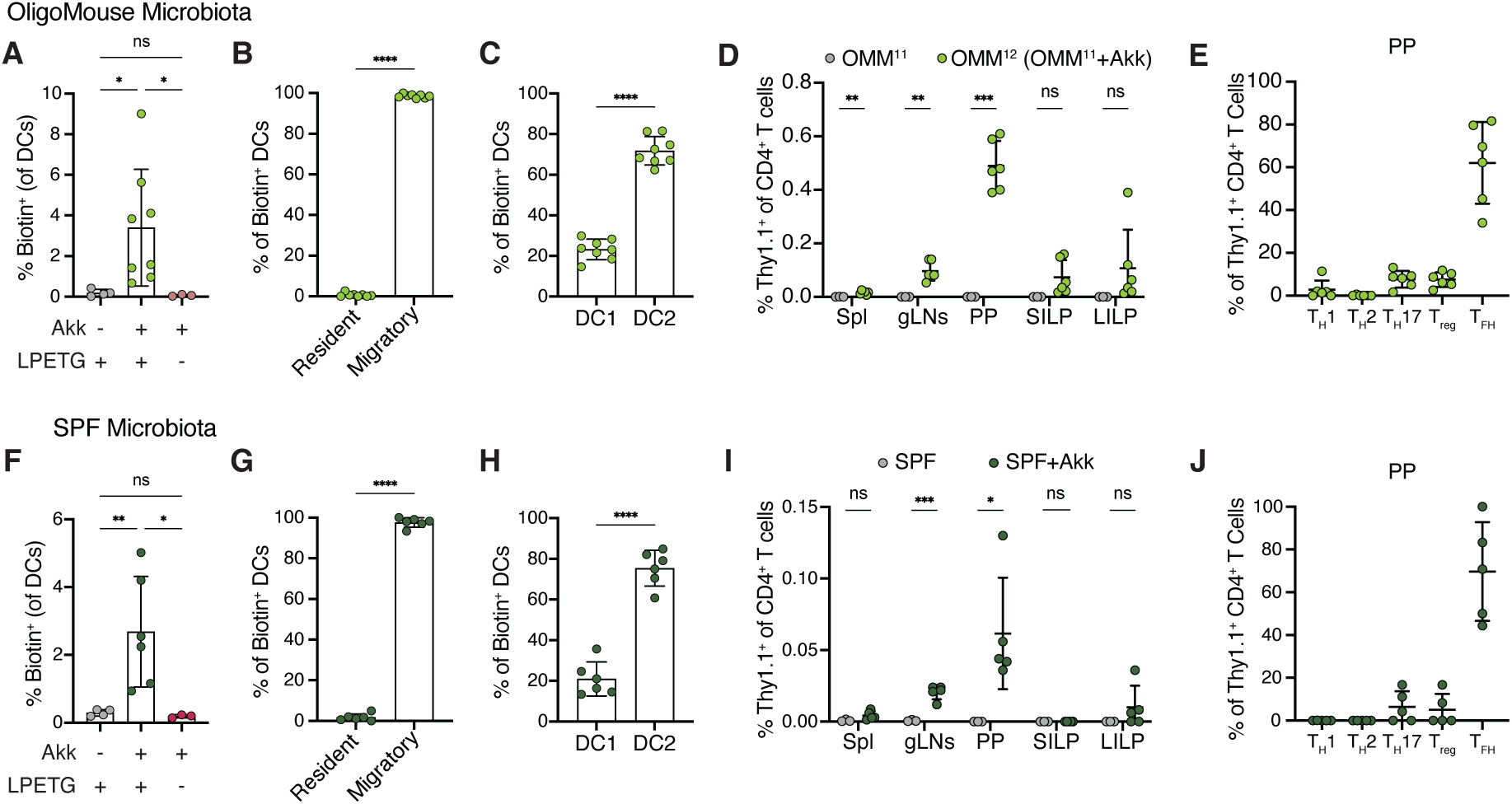
***A. muciniphila*-specific T cells are primed by migratory cDC2s and differentiate into T follicular helper cells in mice with complex microbiota.** For (**A** to **C**), 1×10^6^ *Cd40lg*^SrtAv2^ Am124 T cells were adoptively transferred into OMM^11^ and OMM^12^ (OMM^11^+Akk) mice and LIPSTIC labeling was performed 2.5 days later to identify *A. muciniphila*-presenting cells. (**A**) Percentage of labeled biotin^+^ DCs in gLNs of OMM^11^ and OMM^12^ mice. (**B**) Percentage of migratory and resident cDCs out of total biotin^+^ cDCs from (A). (**C**) Percentage of cDC1s (Xcr1^+^) and cDC2s (Sirpɑ^+^) out of total biotin^+^ cDCs from (A). For (A to C), n = 3 to 8 mice per group, data representative of two independent experiments. For (**D** and **E**), 1×10^4^ Am124 T cells were adoptively transferred into OMM^11^ and OMM^12^ mice. Spleen, gLNs, PPs, SILP, and LILP were harvested 12 days later. (**D**) Frequency of transferred Am124 T cells in tissues of OMM^11^ and OMM^12^ mice. (**E**) Expression of T_H_1 (Tbet^+^ FoxP3^−^), T_H_2 (Gata33^+^ FoxP3^−^), T_H_17 (Rorγt^+^ FoxP3^−^), T_REG_ (FoxP3^+^), and T_FH_ (Bcl6^+^ PD-1^+^ FoxP3^−^) markers in transferred Am124 in the PPs of OMM^12^ mice from (D). For (D and E), n = 3 to 6 mice per group, data representative of two independent experiments. For (**F** to **H**), 1×10^6^ *Cd40lg*^SrtAv2^ Am124 T cells were adoptively transferred into SPF and SPF+Akk mice and LIPSTIC labeling was performed 2.5 days later to identify *A. muciniphila*-presenting cells. (**F**) Percentage of labeled biotin^+^ DCs in gLNs of SPF and SPF+Akk mice. (**G**) Percentage of migratory and resident cDCs out of total biotin^+^ cDCs from (F). (**H**) Percentage of cDC1s (Xcr1^+^) and cDC2s (Sirpɑ^+^) out of total biotin^+^ cDCs from (F). For (F to H), n = 3 to 6 mice per group, data representative of two independent experiments. For (**I** and **J**), 1×10^4^ Am124 T cells were adoptively transferred into SPF and SPF+Akk mice. Spleen, gLNs, PPs, SILP, and LILP were harvested 12 days later. (**I**) Frequency of transferred Am124 T cells in tissues of SPF and SPF+Akk mice. (**J**) Expression of T_H_1 (Tbet^+^ FoxP3^−^), T_H_2 (Gata3^+^ FoxP3^−^), T_H_17 (Rorγt^+^ FoxP3^−^), T_REG_ (FoxP3^+^), and T_FH_ (Bcl6^+^ CXCR5^+^ FoxP3^−^) markers in transferred Am124 in the PPs of SPF+Akk mice from (I). For (I and J), n = 3 to 5 mice per group, data representative of two independent experiments. For all graphs, each symbol represents one mouse and error bars represent mean and standard deviation. P-values were calculated by one-way ANOVA for (A), and (F), and by unpaired T-test for (B to E) and (G to J). Statistical significance denoted as not significant (ns), *P < 0.05, **P < 0.01, ***P < 0.001, ****P < 0.0001.

**Supplementary Figure 4:**
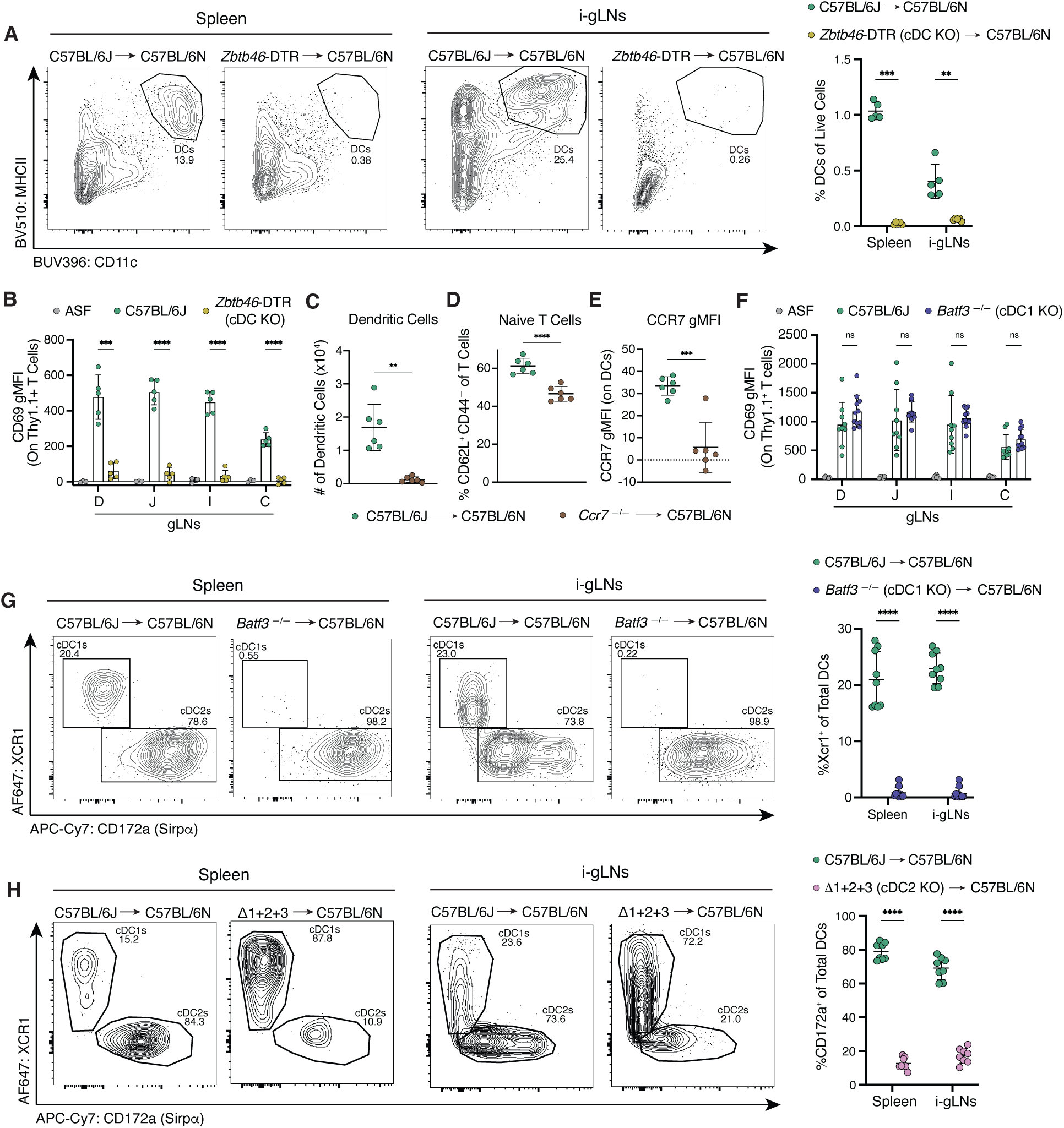
Characterization of ASF+Akk busulfan BMCs. ASF+Akk mice were treated with 25 mg/kg busulfan daily for three days (75 mg/kg total) followed by reconstitution with the indicated bone marrow genotype. 8 to 12 weeks later, 1.5×10^5^Am124 T cells were adoptively transferred into ASF control and ASF+Akk BMC mice. gLNs and spleen were harvested 1.5 days later. (**A**) Representative flow plots (left) and frequency (right) of cDCs in spleen and ileal gLNs of diphtheria-toxin-treated ASF+Akk BMCs reconstituted with C57BL/6J or *Zbtb46*-DTR bone marrow (n = 5 mice per group; data is representative of two independent experiments). (**B**) Expression of CD69 on Am124 T cells in gLNs of ASF control mice or diphtheria-toxin-treated ASF+Akk BMCs reconstituted with C57BL/6J or *Zbtb46*-DTR bone marrow (n = 3 to 5 mice per group, data is representative of two independent experiments). (**C**) Number of DCs in the ileal gLNs of ASF+Akk BMCs reconstituted with C57BL/6J or *Ccr7*^−/–^ bone marrow. (**D**) Frequency of naive (CD62L^+^CD44^−^) T cells in ileal gLNs of ASF+Akk BMCs reconstituted with C57BL/6J or *Ccr7*^−/–^ bone marrow. (**E**) gMFI of CCR7 on DCs in ileal gLNs of ASF+Akk BMCs reconstituted with C57BL/6J or *Ccr7*^−/–^ bone marrow. For (C to E), n = 6 mice per group, data is representative of two independent experiments. (**F**) Expression of CD69 on Am124 T cells 1.5 days post-transfer in gLNs of ASF control mice or ASF+Akk BMCs reconstituted with C57BL/6J or *Batf3*^−/–^ bone marrow (n = 3 to 6 mice per group, per experiment, data is pooled from two independent experiments). (**G**) Representative flow plots (left) and frequency (right) of cDC1s in spleen and ileal gLNs of ASF+Akk BMCs reconstituted with C57BL/6J or *Batf3*^−/–^ bone marrow (n = 4 to 6 mice per group, per experiment, data is pooled from two independent experiments). (**H**) Representative flow plots (left) and frequency (right) of cDC2s in spleen and ileal gLNs from ASF+Akk BMCs reconstituted with C57BL/6J or Δ1+2+3 bone marrow (n = 4 mice per group, per experiment, data is pooled from two independent experiments). For all graphs, each symbol represents one mouse and error bars represent mean and standard deviation. P-values were calculated by unpaired T-test with Welch’s correction for (B), (C), and (F) and unpaired T-test for (A), (D), (E), (G), and (H). Statistical significance denoted as not significant (ns), *P < 0.05, **P < 0.01, ***P < 0.001, ****P < 0.0001.

**Supplementary Figure 5:**
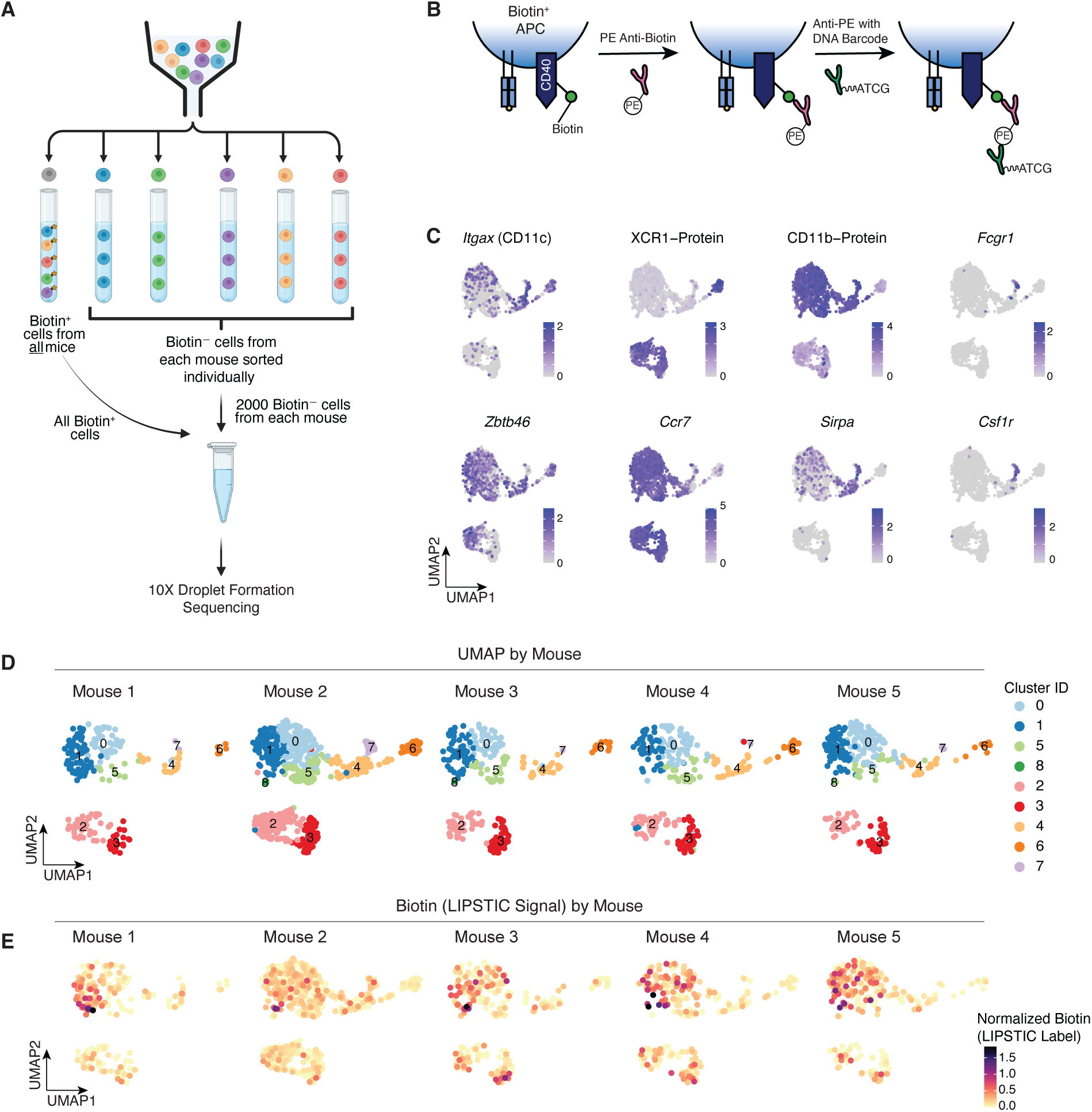
Single cell RNA sequencing of sorted DCs from gLNs. (**A**) Schematic of sorting strategy for single cell sequencing of LIPSTIC labeled DCs. Biotin^+^ DCs from all mice were sorted into one tube. Biotin^−^ cells from each mouse were sorted individually. All biotin^+^ cells were combined with a fraction of the biotin^−^ cells from each mouse. (**B**) Schematic for how biotin (LIPSTIC label) was detected with Totalseq antibodies. Enriched DCs were first incubated with anti-biotin PE antibody so fluorescence could be used for sorting, then cells were incubated with the anti-PE TotalSeq antibody with a DNA barcode. (**C**) Feature plots showing the expression of key marker genes that were used to define cDC1s, cDC2s and migratory DCs clusters. Xcr1 and CD11b were detected at the protein level using TotalSeq antibodies. All other genes were detected at the RNA level. (**D**) Uniform manifold approximation and projection (UMAP) plot of sorted DCs from gLNs in individual mice. All clusters contained cells from all mice. (**E**) Log normalized counts of LIPSTIC signal in individual mice. LIPSTIC signal detected via anti-PE hashtag antibody as outlined in (B).

**Supplementary Figure 6:**
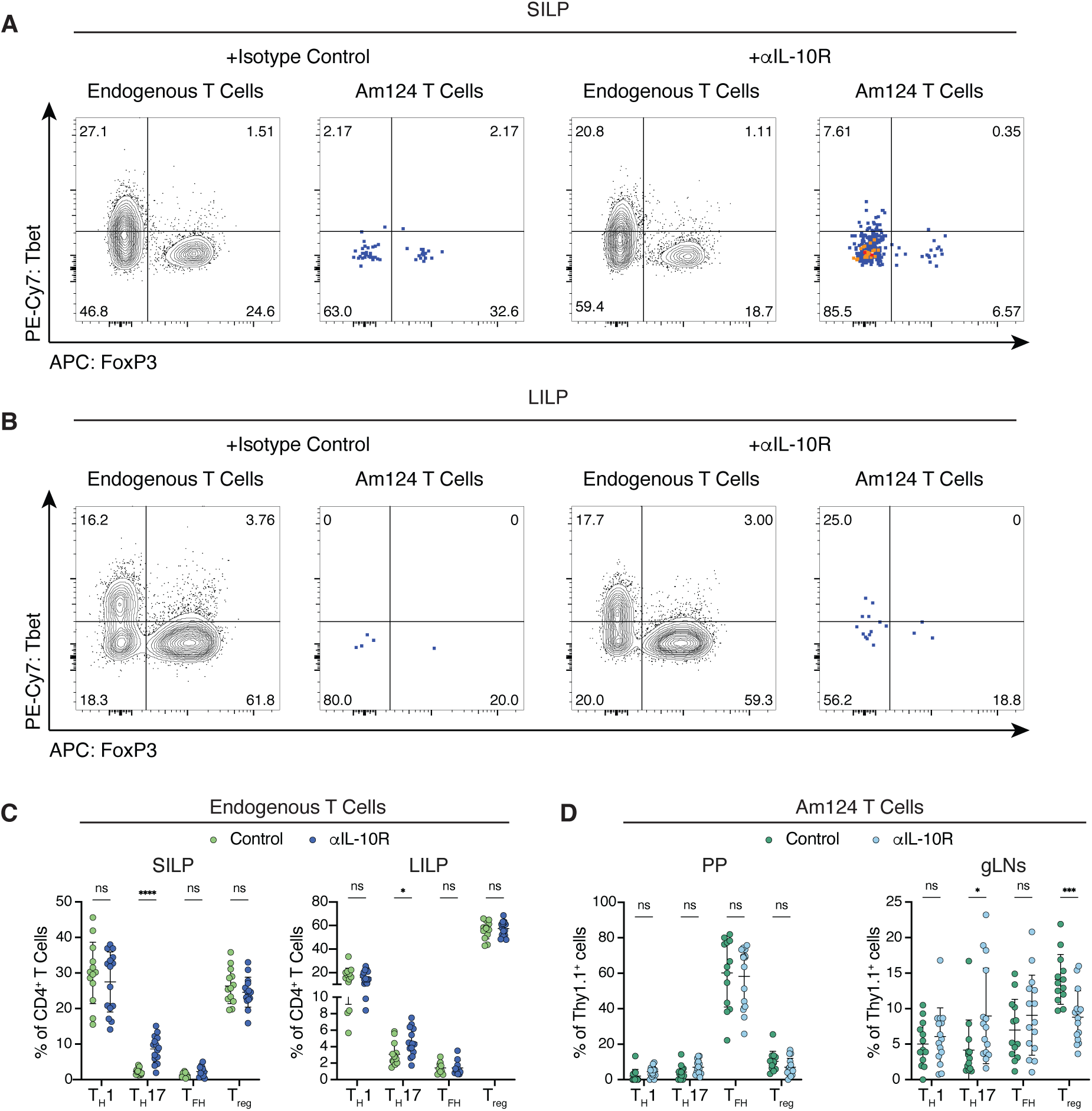
Additional characterization of endogenous and Am124 T cells in αIL-10R-treated mice. (**A** and **B**) Representative flow plots showing expression of the T_H_1 marker Tbet by endogenous and transferred T cells in the (**A**) SILP and (**B**) LILP of control mice and ɑIL-10R-treated mice. (**C**) Frequency of endogenous CD4^+^ T cells expressing T_H_1 (Tbet^+^ FoxP3^−^), T_H_17 (Rorγt^+^ FoxP3^−^), T_FH_ (Bcl6^+^ PD-1^+^) and T_REG_ (FoxP3^+^) markers in the SILP (left) and LILP (right) of control and ɑIL-10R-treated mice. (**D**) Frequency of transferred Am124 T cells expressing T_H_1 (Tbet^+^ FoxP3^−^), T_H_17 (Rorγt^+^ FoxP3^−^), T_FH_ (Bcl6^+^ PD-1^+^) and T_REG_ (FoxP3^+^) markers in the PPs (left), and gLNs (right) of control and ɑIL-10R-treated mice. For all graphs, each symbol represents one mouse and the error bars represent mean and standard deviation; n = 4 to 6 mice per group, per experiment, data is pooled from three independent experiments. P-values were calculated by unpaired T-test. Statistical significance denoted as not significant (ns), *P < 0.05, **P < 0.01, ***P < 0.001, ****P < 0.0001.

**Supplementary Figure 7:**
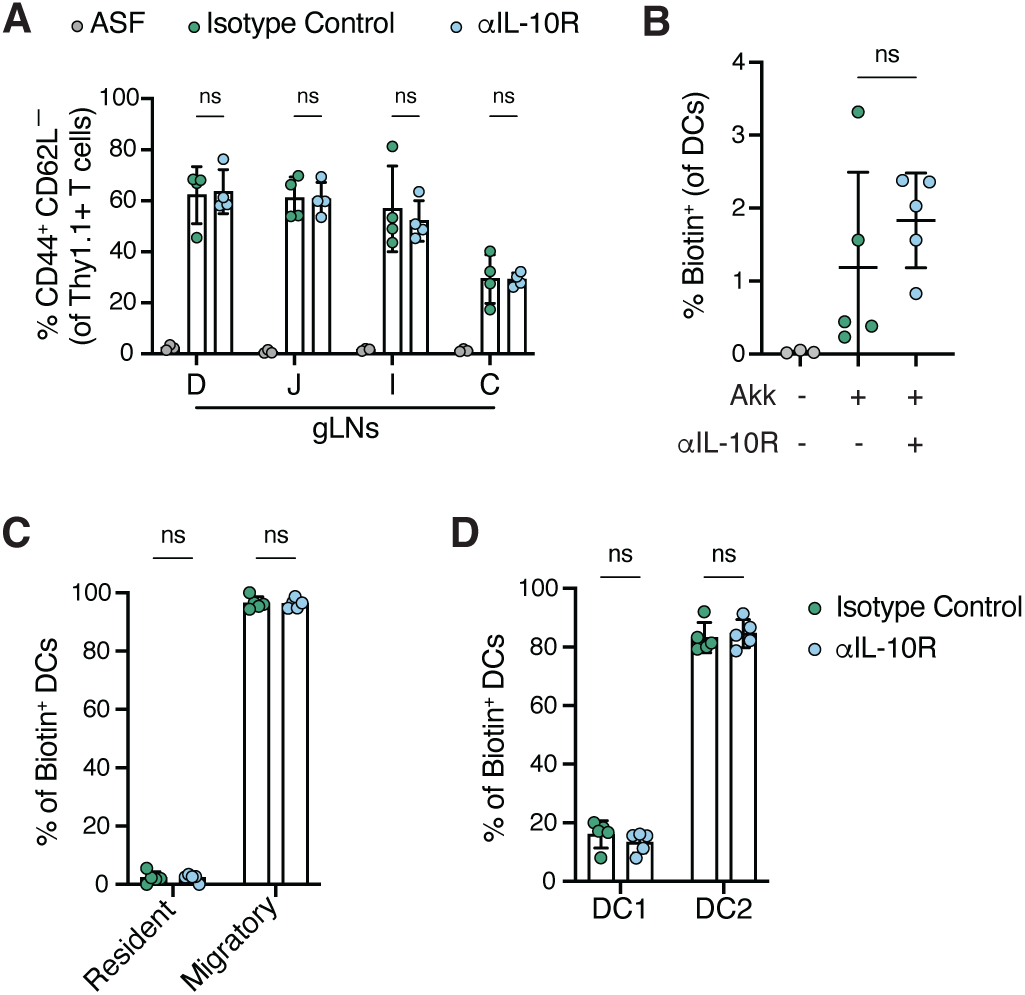
αIL-10R treatment does not alter *A. muciniphila*-specific T cell priming in the gLNs or the identity of presenting APC. (**A**) 2.5×10^5^ Am124 T cells were adoptively transferred into ASF (untreated) and ASF+Akk mice 3 days after injection of 1 mg ɑIL-10R or isotype control antibodies. gLNs were harvested 2.5 days post-transfer. Percentage of activated (CD44^+^ CD62L^−^) Am124 T cells in gLNs of ASF and isotype- or ɑIL-10R-treated ASF+Akk mice (n = 3 to 4 per group, data is representative of two independent experiments). For (**B** to **D**), 1x×0^6^ *Cd40lg*^SrtAv2^ Am124 T cells were adoptively transferred into ASF (untreated) and ASF+Akk mice 3 days after injection of 1 mg ɑIL-10R or isotype control antibodies. LIPSTIC labeling was performed 2.5 days later to identify *A. muciniphila*-presenting cells. (**B**) Percentage of LIPSTIC-labeled biotin^+^ DCs in gLNs of ASF and isotype- or ɑIL-10R-treated ASF+Akk mice. (**C**) Percentage of migratory and resident cDCs out of total biotin^+^ cDCs in ASF+Akk mice from (B). (**D**) Percentage of cDC1s (Xcr1^+^) and cDC2s (Sirpɑ^+^) cDCs out of total biotin^+^ cDCs from ASF+Akk mice in (B). For (B to D), n = 3 to 5 mice per group, data is representative of three independent experiments. For all graphs, each symbol represents one mouse and error bars represent mean and standard deviation. P-values were calculated by unpaired T-test for (A), (C), and (D), and by one-way ANOVA for (B). Statistical significance denoted as not significant (ns), *P < 0.05, **P < 0.01, ***P < 0.001, ****P < 0.0001.

**Supplementary Figure 8:**
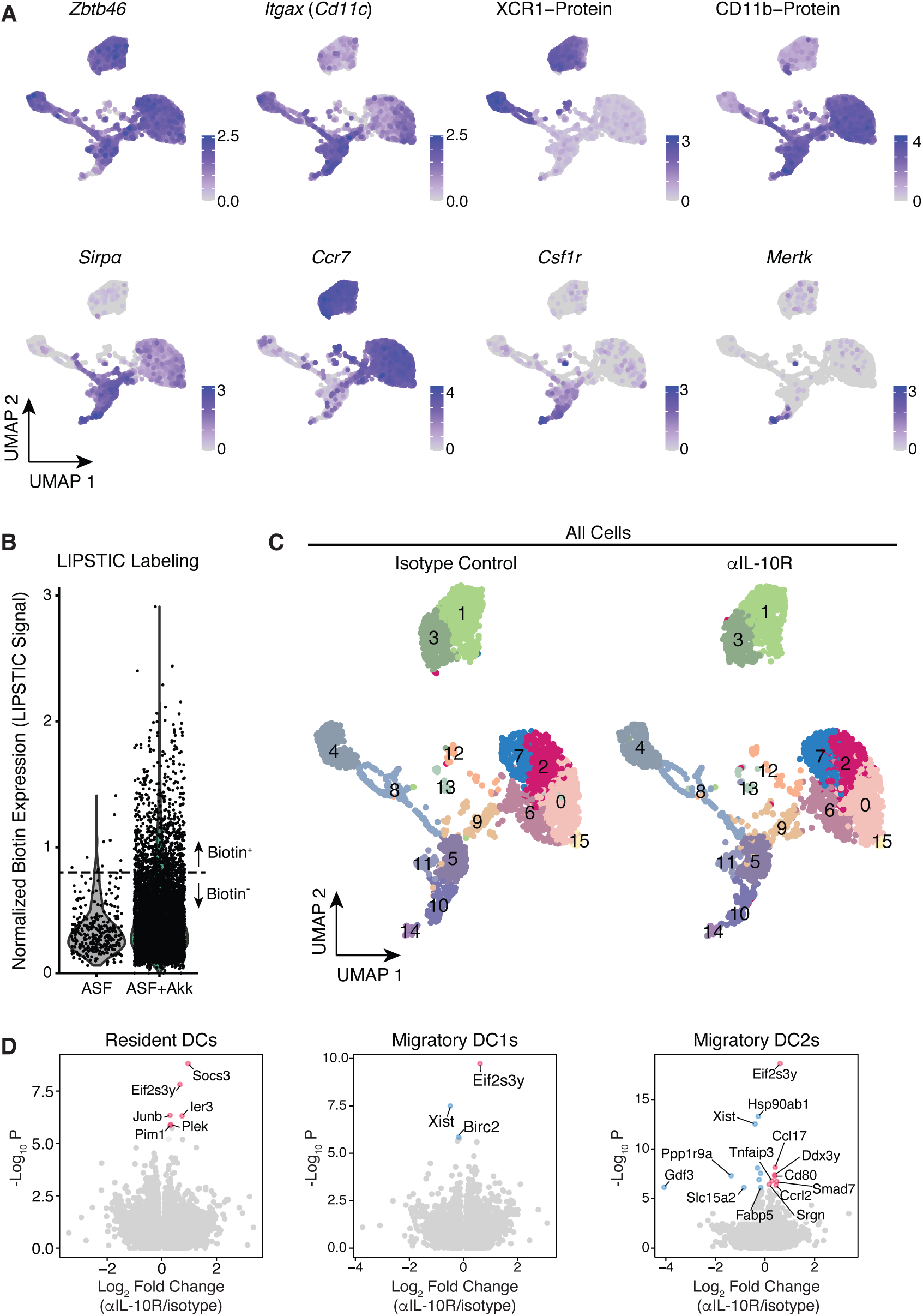
Single cell RNA sequencing of sorted cDCs from gLNs of ASF+Akk mice treated with ɑIL-10R or isotype control antibody. (**A**) Feature plots showing the expression of key marker genes that were used to define cDC1, cDC2 and migratory cDCs clusters. Xcr1 and CD11b were detected at the protein level using TotalSeq antibodies. All other genes were detected at the RNA level. (**B**) Log normalized counts of LIPSTIC signal in cells from ASF and isotype- or ɑIL-10R-treated ASF+Akk mice detected via anti-PE hashtag antibody. Dashed line at 0.8 is the cutoff used in Fig. 4 for biotin^+^ cells. (**C**) Uniform manifold approximation and projection (UMAP) plot of sorted DCs from gLNs separated by treatment group. (**D**) Volcano plots showing differentially expressed genes between mice treated with ɑIL- 10R and isotype control antibody for resident cDCs (left), migratory cDC1s (middle), and migratory cDC2s (right). Statistically significant genes are colored in red if they are enriched in ɑIL-10R-treated mice or blue if they are enriched in isotype control mice.

**Supplementary Figure 9:**
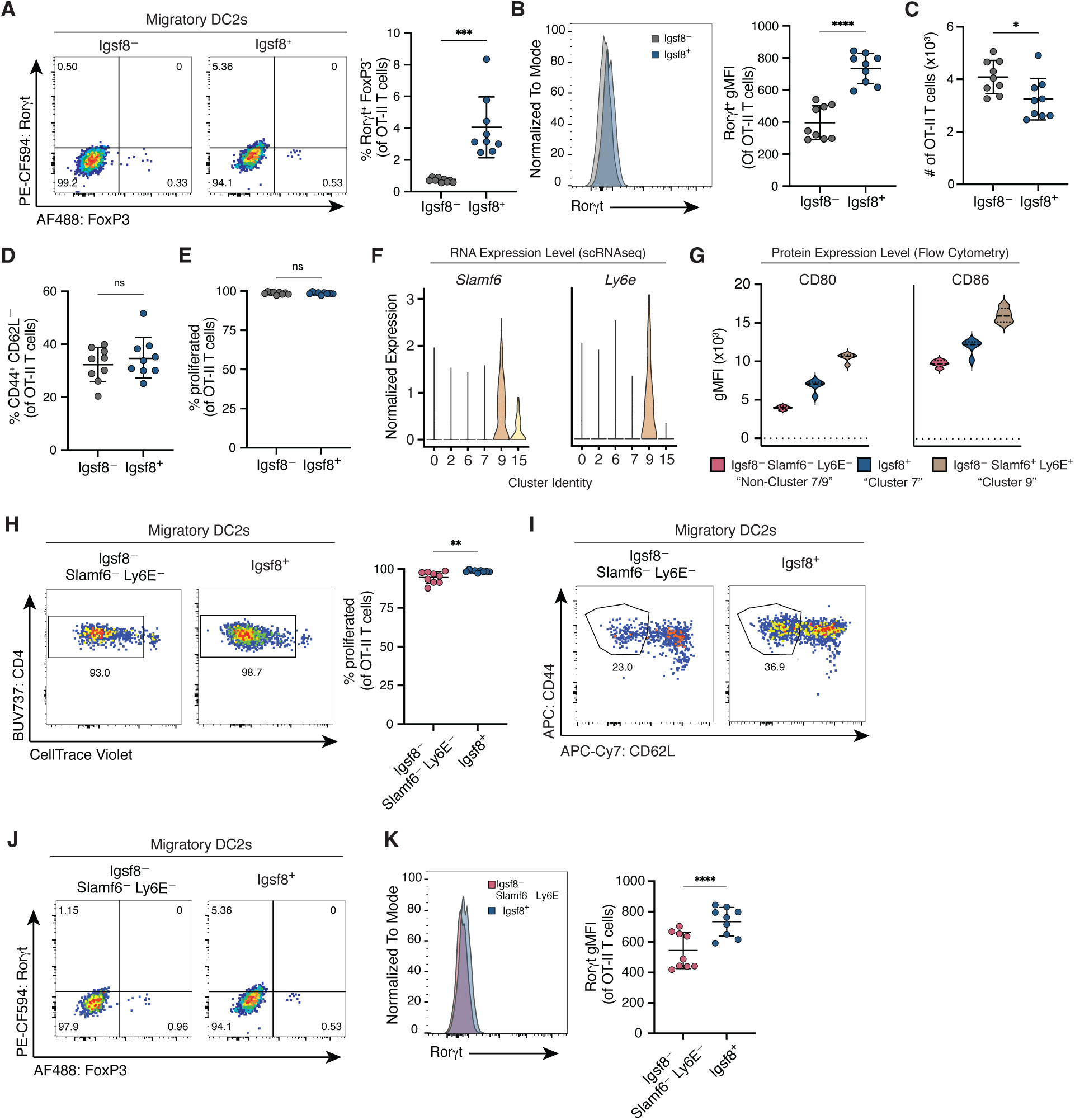
Distinct migratory cDC2 subpopulations differentially promote T cell activation and T_H_17 differentiation *in vitro*. For (**A** to **E**, and **H** to **K**), naive, CTV-labeled OT-II CD4^+^ T cells were co-cultured in the presence of exogenous OT-II peptide for 96 hours with the indicated migratory cDC2 populations sorted from the gLNs of untreated ASF+Akk mice. (**A**) Representative flow plots (left) and frequency (right) of Rorγt^+^ FoxP3^−^ OT-II T cells after co-culture with Igsf8^−^ or Igsf8^+^ migratory cDC2s. (**B**) Representative histograms (left) and quantification (right) of Rorγt expression in OT-II T cells after co-culture with Igsf8^−^ or Igsf8^+^ migratory cDC2s. (**C**) Total number of OT-II T cells after co-culture with Igsf8^−^ or Igsf8^+^ migratory cDC2s. (**D**) Frequency of activated (CD44^+^ CD62L^−^) OT-II cells after co-culture with Igsf8^−^ or Igsf8^+^ migratory cDC2s. (**E**) Frequency of proliferated OT-II cells after co-culture with Igsf8^−^ or Igsf8^+^ migratory cDC2s. (**F**) Expression of the indicated genes at the RNA level in bulk migratory cDC2 clusters from scRNAseq experiment in Fig. 5. (**G**) Expression of the indicated genes at the protein level in Igsf8^−^ Slamf6^−^ Ly6E^−^ (“non-cluster 7/9”) Igsf8^+^ (“cluster 7) and Igsf8^−^ Slamf6^+^ Ly6E^+^ (“cluster 9”) migratory cDC2s. Solid and dashed lines indicate median and quartiles. n = 4 to 5 mice per experiment, data representative of two independent experiments. (**H**) Representative flow plots (left) and frequency (right) of proliferated OT-II T cells after co-culture with Igsf8^−^ Slamf6^−^ Ly6E^−^ or Igsf8^+^ migratory cDC2s. (**I**) Representative flow plots of activated (CD44^+^ CD62L^−^) OT-II T cells after co-culture with Igsf8^−^ Slamf6^−^ Ly6E^−^ or Igsf8^+^ migratory cDC2s. (**J**) Representative flow plots of Rorγt^+^ FoxP3^−^ OT-II T cells after co-culture with Igsf8^−^ Slamf6^−^ Ly6E^−^ or Igsf8^+^ migratory cDC2s. (**K**) Representative histograms (left) and quantification (right) of Rorγt expression in OT-II T cells after co-culture with Igsf8^−^ Slamf6^−^ Ly6E^−^ or Igsf8^+^ migratory cDC2s. For (A to E, and H to K), n = 4 to 5 mice per experiment; data pooled from two independent experiments. P-values were calculated via paired T-test. For all graphs, each symbol represents one mouse and error bars represent mean and standard deviation. Statistical significance denoted as not significant (ns), *P < 0.05, **P < 0.01, ***P < 0.001, ****P < 0.0001.

**Supplementary Figure 10:**
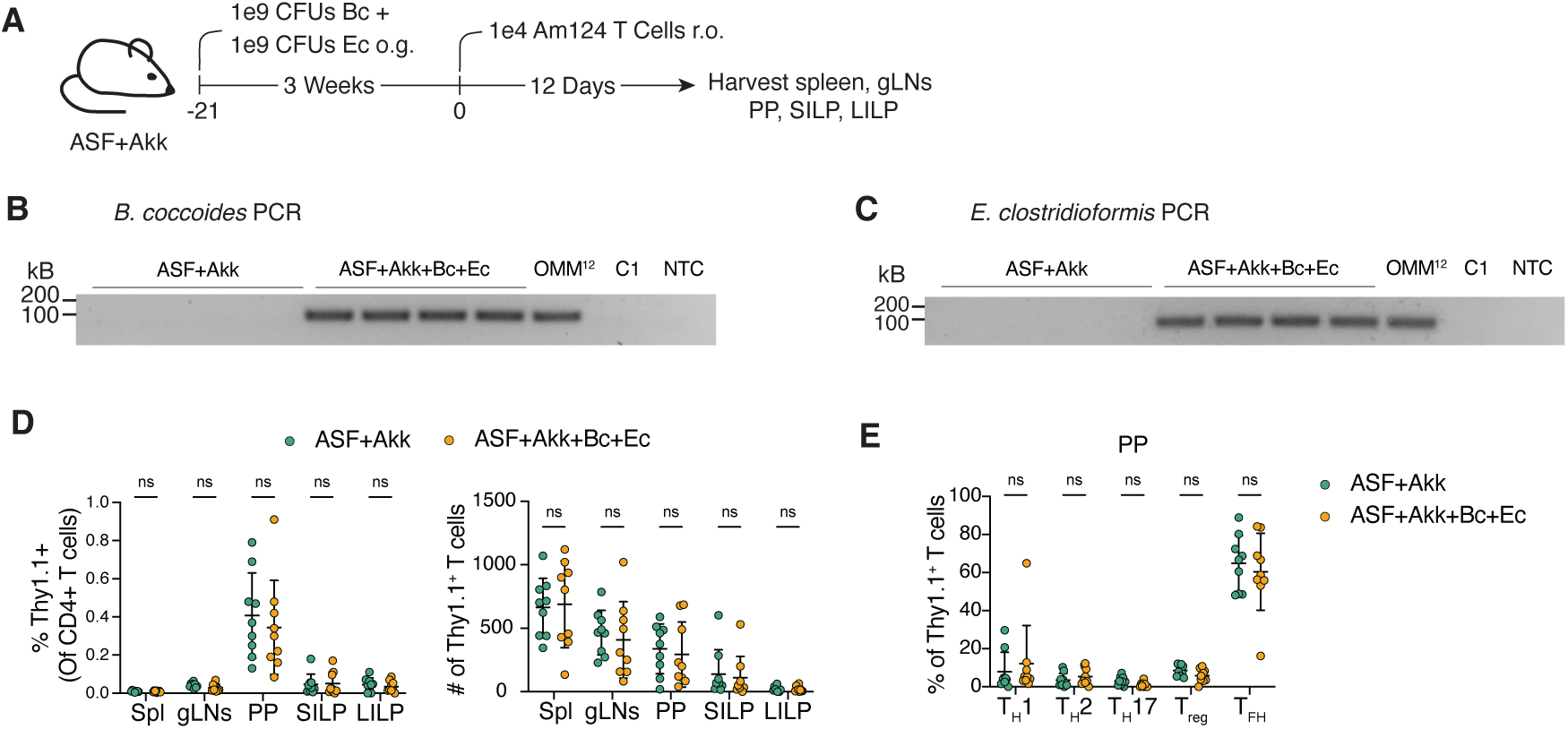
Colonization with *B. coccoides* and *E. clostridioformis* does not alter the differentiation of *A. muciniphila*-specific T cells. (**A**) Schematic representation of experimental timeline. 1×10^9^ CFUs of *B. coccoides* (Bc) and 1×10^9^ CFUs of *E. clostridioformis* (Ec) were gavaged into ASF+Akk mice to make ASF+Akk+Bc+Ec mice. Three weeks post-gavage, 1×10^4^ Am124 T cells were adoptively transferred into ASF+Akk and ASF+Akk+Bc+Ec mice. Spleen, gLNs, PPs, SILP, and LILP were harvested 12 days post-transfer. (**B** and **C**) Agarose gel of *B. coccoides*- (**B**) and *E. clostridioformi*s- (**C**) specific amplicons from fecal DNA of indicated mice. Fecal DNA from OMM^12^ mice is used as a positive control. Two negative controls are included: a contamination control for the fecal prep (C1) and a non-template control (NTC). (**D**) Frequency (left) and number (right) of transferred Am124 T cells in tissues of ASF+Akk mice and ASF+Akk+Bc+Ec 12 days post-transfer. (**E**) Frequency of transferred Am124 T cells expressing T_H_1 (Tbet^+^ FoxP3^−^), T_H_2 (Gata3^+^ FoxP3^−^), T_H_17 (Rorγt^+^ FoxP3^−^), T_FH_ (Bcl6^+^ PD-1^+^) and T_REG_ (FoxP3^+^) markers in the PPs tissues of ASF+Akk mice and ASF+Akk+Bc+Ec. For (D and E), n = 4 to 5 mice per group, per experiment, data is pooled from two independent experiments. Each symbol represents one mouse and error bars represent mean and standard deviation. P-values were calculated using unpaired T-test. Statistical significance denoted as not significant (ns), *P < 0.05, **P < 0.01, ***P < 0.001, ****P < 0.0001.

**Table S1. (separate file)**

Primers and probes used for detecting bacterial species

**Table S2. (separate file)**

Antibodies and dyes used for flow cytometry

**Table S3. (separate file)**

Genes negative and positively correlated with biotin signal in migratory cDC2s in Fig. 3

**Table S4. (separate file)**

sc2markers output for cluster 7 migratory cDC2s in Fig. 5

**Table S5. (separate file)**

sc2markers output for cluster 9 migratory cDC2s in fig. S9

